# Differential Impacts on Host Transcription by ROP and GRA Effectors from the Intracellular Parasite *Toxoplasma gondii*

**DOI:** 10.1101/2020.02.04.934570

**Authors:** Suchita Rastogi, Yuan Xue, Stephen R. Quake, John C. Boothroyd

**Affiliations:** Department of Microbiology and Immunology, Stanford University School of Medicine, Stanford, CA, USA; Department of Bioengineering, Stanford University, Stanford, CA, USA; Department of Applied Physics, Stanford University, Stanford, CA, USA; Chan Zuckerberg Biohub, San Francisco, CA, USA

## Abstract

The intracellular parasite *Toxoplasma gondii* employs a vast array of effector proteins from the rhoptry and dense granule organelles to modulate host cell biology; these effectors are known as ROPs and GRAs, respectively. To examine the individual impacts of ROPs and GRAs on host gene expression, we developed a robust, novel protocol to enrich for ultra-pure populations of a naturally occurring and reproducible population of host cells called uninfected-injected (U-I) cells, which *Toxoplasma* injects with ROPs but subsequently fails to invade. We then performed single cell transcriptomic analysis at 1-3 hours post-infection on U-I cells (as well as on uninfected and infected controls) arising from infection with either wild type parasites or parasites lacking the MYR1 protein, which is required for soluble GRAs to cross the parasitophorous vacuole membrane (PVM) and reach the host cell cytosol. Based on comparisons of infected and U-I cells, the host’s earliest response to infection appears to be driven primarily by the injected ROPs, which appear to induce immune and cellular stress pathways. These ROP-dependent pro-inflammatory signatures appear to be counteracted by at least some of the MYR1-dependent GRAs and may be enhanced by the MYR-independent GRAs, (which are found embedded within the PVM). Finally, signatures detected in uninfected bystander cells from the infected monolayers suggests that MYR1-dependent paracrine effects also counteract inflammatory ROP-dependent processes.

**IMPORTANCE:** This work performs the first transcriptomic analysis of U-I cells, captures the earliest stage of a host cell’s interaction with *Toxoplasma gondii*, and dissects the effects of individual classes of parasite effectors on host cell biology.

## Introduction

The obligate intracellular parasite *Toxoplasma gondii* parasitizes a wide range of avian and mammalian organisms, including humans (1). During the acute phase of infection, this unicellular eukaryote rapidly expands within host tissues by penetrating host cells, establishing and replicating within an intracellular parasitophorous vacuole (PV), and simultaneously avoiding clearance by the host immune system (reviewed in (2)). To orchestrate these events, *Toxoplasma* employs a vast repertoire of effector proteins housed primarily in two secretory organelles, the rhoptries and dense granules (Fig. 1A). The rhoptry organelle effectors (ROPs) are secreted by the parasite into the host cytosol during or immediately prior to invasion by an as yet undefined mechanism (reviewed in (3)). The handful of ROPs that have been characterized to date include virulence factors that disrupt immune clearance mechanisms (4–7), remodel the host’s cortical actin cytoskeleton at the point of parasite penetration (8, 9), and co-opt the STAT3 and STAT6 pathways (10–12). In contrast, the dense granule effectors (GRAs) are thought to be secreted later during the invasion process and mostly after the parasite has invaded the host cell (reviewed in (3)). Many GRAs remain in the PV lumen or PV membrane (PVM) (13–17), but others traverse the PVM to reach the host cytosol and often proceed to the host nucleus (18–24). The translocation of this latter class of GRAs across the PVM is dependent on a group of PVM proteins called the MYR complex, i.e., MYR1, MYR2, and MYR3, which are so-named because they are required for parasite-dependent host c-<u>My</u>c regulation (25–27). Host signaling pathways modulated by the PVM-embedded, MYR-independent GRAs (MIGs) include the nuclear factor kappa light chain enhancer of activated B cells pathway. Host pathways modulated by MYR-dependent GRAs (MDGs) include the interferon gamma (IFN-gamma), mitogen activated protein kinase, and p53 pathways, as well as the cyclin E regulatory complex (22, 23, 28).

**Figure 1.**
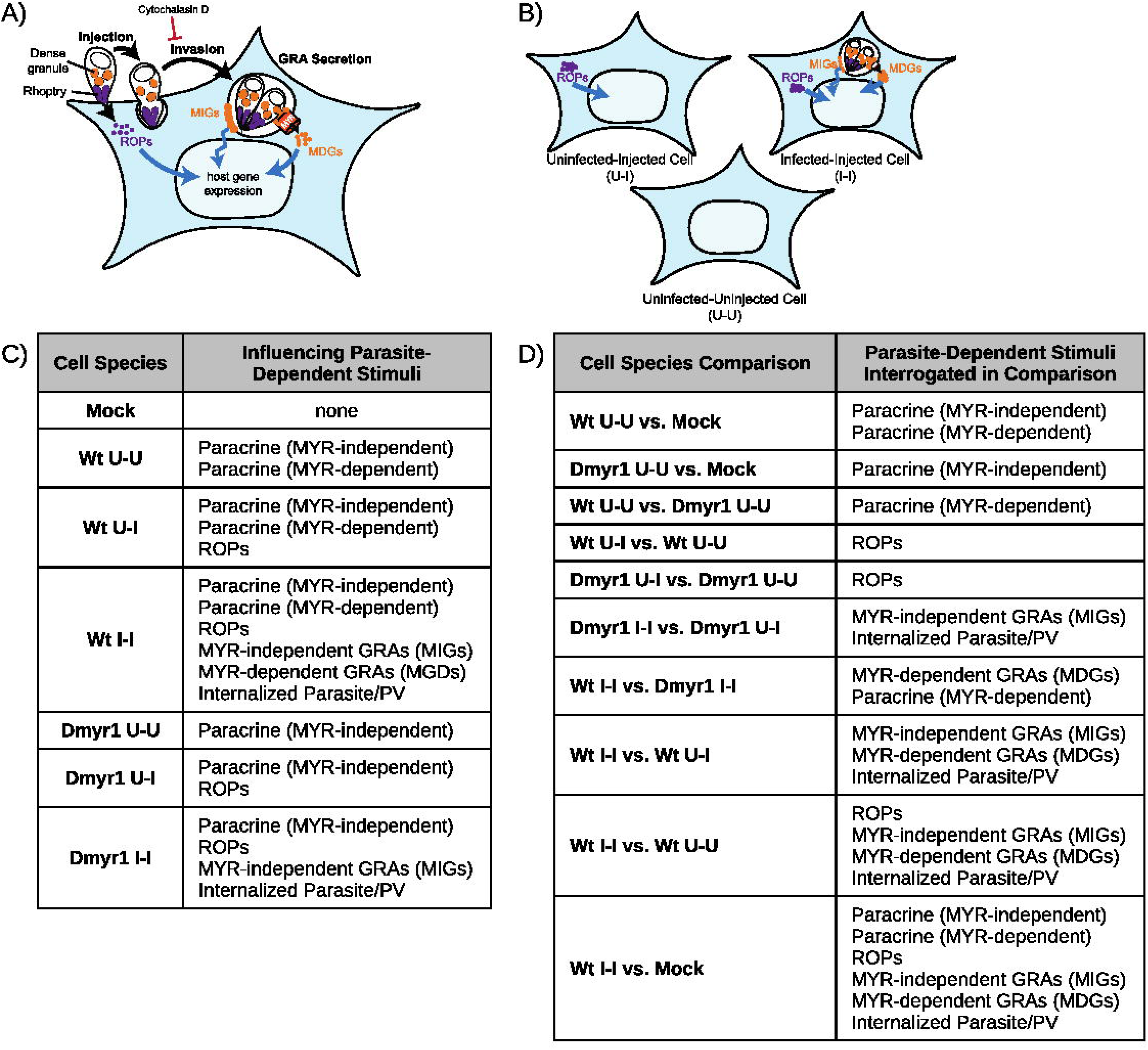
Key experimental conditions and comparisons. **(A)** Simplified illustration of effector secretion during early tachyzoite infection. Blue curved arrows = modulation of host transcription, black curved arrows = transitions through stages of lytic cycle, blue jagged arrow = modulation without translocation to the nucleus, ROPs = rhoptry proteins, MIGs = MYR-independent dense granule proteins, MDGs = MYR-dependent dense granule proteins, MYR = putative translocation required for MDG penetration of host cytosol. **(B)** As a result of the events in (A), infected host cell monolayers produce infected-injected (I-I) cells when tachyzoite invasion proceeds as usual, uninfected-uninjected (U-U) bystander cells, and uninfected-injected (U-I) cells from aborted invasion after effector injection. **(C)** Parasite-dependent stimuli (i.e., effectors, paracrine factors) that influence each of seven cell species collected for RNA sequencing, according to the current model of effector secretion for *Toxoplasma* tachyzoites. Wt and Dmyr1 designate cells originating from host cell monolayers infected with wild type and D*myr1* parasites, respectively; Mock indicates cells from a mock-infected monolayer. **(D)** Parasite-dependent stimuli that explain the differences in expression trends between key pairs of collected cell species. PV = parasitophorous vacuole.

The collective host response to ROPs, MDGs, MIGs, and other perturbations during infection with *Toxoplasma gondii* have been well-documented in transcriptomic studies that compare infected and mock-infected cells; however, the contribution of individual parasite effector compartments to the overall picture is more poorly defined, particularly because many parasite effectors are introduced during the earliest stages of infection in very narrow time intervals. Specifically, invasion is a rapid process that takes approximately 40 seconds (29), and deployment of the ROPs likely occurs during the first third of invasion in one shot (30). In contrast, MDGs have been detected by immunofluorescence assays in the nuclei of parasitized host cells at ∼3 hours post-invasion at the earliest (19), which suggests that they likely modulate host transcription much later than ROPs. Parasite mutants that lack a functional MYR complex have helped separate the impact of MDGs from those of other parasite effectors (25, 26, 31), but the specific impact on host transcription by ROPs and MIGs has yet to be determined. The rhoptry organelle’s contribution is of particular interest given that ROP injection is essential to parasite invasion, survival, and virulence, and that the functions of most ROPs are unknown.

Here we document the impact of specific classes of parasite effectors, including for the first time ROPs, by leveraging a rare population of host cells that parasites inject with ROPs but subsequently fail to invade (32). These <u>u</u>ninfected-injected (U-I) host cells (Fig. 1B) do not arise as a result of parasite-derived exosomes delivering effectors to host cells (32) or due to impending host cell death (Fig. 2B). They arise spontaneously in tissue culture during *Toxoplasma* infection and may also arise *in vivo* in the brains of mice chronically infected with *Toxoplasma* (32). To interrogate the host response specifically to effectors injected before invasion, we developed a novel pipeline to purify and perform single cell RNA sequencing (scRNA-seq) of U-I cells, as well as of infected and uninfected controls from the same host cell monolayer. To resolve the impact of effectors released during vs. after invasion, we also analyzed U-I cells, infected cells, and uninfected cells from host cell monolayers infected with parasites lacking MYR1, a component of the complex required for translocation of MDGs into the host cytosol (25, 26). Our findings reveal new insight into the impact of individual parasite effector compartments on the biology of *Toxoplasma*-infected host cells.

**Figure 2.**
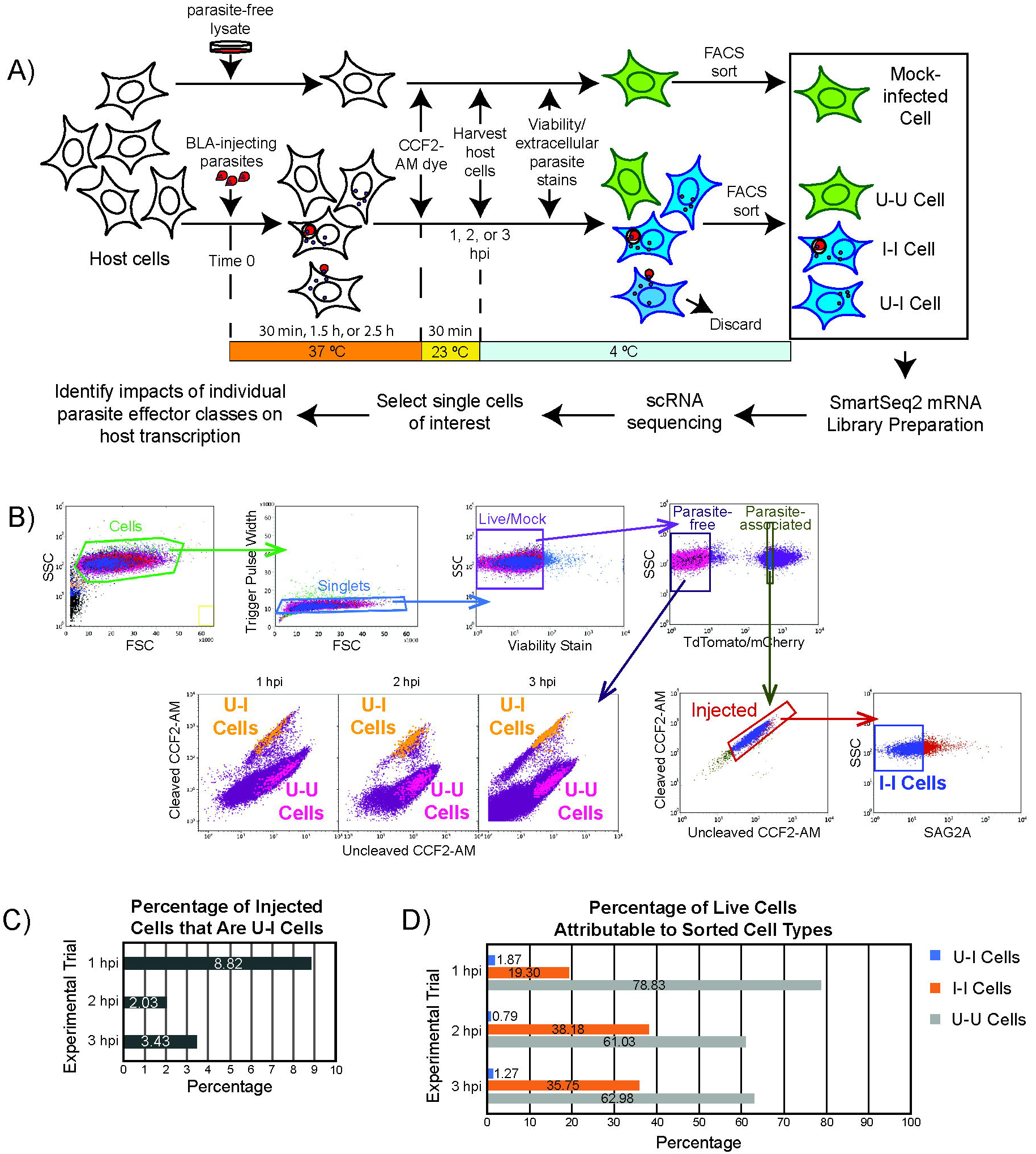
Experimental pipeline. **(A)** Pipeline to collect and analyze experimental conditions for single cell RNA sequencing. BLA = toxofilin-beta-lactamase ROP fusion protein, CCF2-AM = indicator dye that reveals injected (blue) vs. uninjected (green) cells, U-U = uninfected-uninjected bystander host cell, U-I = uninfected-injected host cell, and I-I = infected-injected host cell. **(B)** FACS gating strategy to obtain mock-infected, U-U, I-I, and U-I cells. For cells originating from the parasite-free gate, the distribution of sorted U-I (orange) and U-U (pink) cells from each of the 1 hpi, 2 hpi, and 3 hpi time points is shown. **(C)** Proportion of injected (U-I + I-I) cells that are U-I. **(D)** Abundances of U-U, I-I, and U-I cells during FACS, expressed as percentages of cells from the Live/Mock gate from which they were sorted.

## Results

### A Novel FACS-Based Single Cell RNA Sequencing Pipeline Captures Transcriptomic Signatures of Individual Effector Compartments

According to the current model of parasite effector secretion (Fig. 1A), the host response to an infection with *Toxoplasma* tachyzoites can be attributed to 5 parasite-dependent stimuli: 1) the secretion of paracrine effectors into the extracellular milieu; 2) injection of ROPs; 3) activity of MIGs; 4) secretion of MDGs; and 5) parasite invasion and establishment of the PV. To resolve the individual impacts of each of these classes of stimulus, we transcriptomically profiled infected, uninfected, and U-I host cells from monolayers exposed to parasites. To this end, we devised a novel pipeline in which: 1) infections were designed to generate a heterogeneous pool of host cells each impinged upon by one of several combinations of parasite-dependent stimuli; 2) a fluorescence activated cell sorting (FACS)-based protocol was employed to separate host cells of interest from infected monolayers based on the parasite-dependent stimuli by which they were affected; and 3) the host cell transcriptomes were profiled by full-length scRNA-seq. Given the novelty and complexity of this pipeline, before we delineate our biological findings, here we present an account of our strategy and technical findings during pipeline development.

### FACS-Based Isolation of Host Cells for Single Cell RNA Sequencing Purifies

### Uninfected-Injected Cells

To generate host cells impacted by various combinations of parasite-dependent stimuli for downstream RNA-seq, we subjected the host cells to infection with either wild type RH (type I) strain parasites, mutant *Δmyr1* parasites lacking the MYR1 protein (constructed from an RH *△myr1* mCherry parental strain (25); Fig. S1)), or a parasite-free cell lysate (i.e., mock infection) from the feeder human foreskin fibroblast (HFF) cells used to maintain both parasite lines. From here on, we designate host cells that arose from the wild type, *Δmyr1*, and mock infections as Wt, Dmyr1, or Mock cells, respectively. All parasites were engineered to express the ROP fusion protein toxofilin-beta-lactamase (Tfn-BLA) as well as a constitutive red fluorescent protein. In addition, we chose 10 T1/2 mouse fibroblasts as the preferred host cell type over conventionally used HFFs, which we found unsuitable for single cell sorts due to their propensity to remain clumped in FACS buffer.

As each true infection results in a heterogeneous monolayer containing U-I cells, infected cells (designated I-I for being infected and injected), and uninfected cells (denoted U-U for being <u>u</u>ninfected and <u>u</u>ninjected; Fig. 1B), the infection conditions yielded seven key species of host cell: Wt U-I, Wt I-I, Wt U-U, Dmyr1 U-I, Dmyr1 I-I, Dmyr1 U-U, and Mock. Each of these species was expected to be influenced by a specific combination of parasite-dependent stimuli (Fig. 1C). By comparing the transcriptomes of key pairs of these seven cell species (see Materials and Methods, Data Availability), we expected to uncover the impact of previously unexamined parasite-dependent stimuli, e.g., ROPs (summarized in Fig. 1D). Of note, these experimental conditions enabled transcriptomic assessment of paracrine effects during *Toxoplasma* infection (by comparison of U-U vs. mock-infected cells) and generated pure I-I and U-I cell populations at extremely short infection durations, an improvement over traditional methods that rely on high multiplicities of infection and long infection times to distinguish infected vs. uninfected transcriptomic signatures.

To capture individual cells from each cell species, we devised a FACS-based protocol that purified each species from the host cell monolayers (Fig. 2A). The pipeline employed the toxofilin-beta-lactamase (Tfn-BLA) assay (8), in which the (mock)-infected host cell monolayers were stained for 30 minutes with CCF2-AM, a reporter dye that is taken up by live host cells and shifts from green to blue fluorescence when cleaved by beta-lactamase. CCF2-AM staining occurred at 30, 90, or 150 minutes post-infection, to yield cells at 1, 2, or 3 hours post-infection (hpi). In addition, CCF2-AM staining was performed at room temperature (23 °C) to preserve the integrity of the CCF2-AM dye, which decomposes at 37 °C. U-I cells were then FACS sorted based on their blue (cleaved) CCF2-AM fluorescence and their absent red fluorescence (since they lacked internalized parasites). In addition, I-I cells were sorted for their double positive blue (from cleaved CCF2-AM) and red (from internalized parasites) fluorescence, and U-U and Mock cells were sorted for their green (cleaved) CCF2-AM fluorescence. Single cells and bulk populations of 50-100 cells of each cell species were collected.

To ensure confidence in the identity of the sorted cells and to limit the number of parasites per I-I cell to one, we gated stringently during FACS, particularly for the “U-I cells” gate and “parasite-associated” gate (which included I-I cells; Fig. 2B). We also subjected all cells to extensive washes before FACS sorting, as debris from the human feeder cells used to culture the parasites contaminates the U-I gate (Fig. S2), likely due to retention of both CCF2-AM dye and the parasite fusion protein Tfn-BLA. Of note, the debris posed significant challenges to downstream bioinformatic analyses of bulk U-I cells, but they registered as single cells in the scRNA-seq pipeline and were automatically discarded during quality control due to their high percentage of human reads. At a multiplicity of infection of ∼6, U-I cells constituted ∼8.8% of the injected (i.e., U-I + I-I) cell population at 1 hpi, ∼2.0% at 2 hpi, and ∼3.4% at 3 hpi (Fig. 2C). Abundances of sorted U-I, I-I, and U-U cells relative to all sorted cells are depicted in Fig. 2D.

To validate U-I cells as appropriate models for host responses to injected parasite effectors, we also limited the possibility of U-I cells arising by mechanisms other than aborted invasion events. One such mechanism is host immune clearance of internalized parasites. However, this route is an unlikely source of U-I cells in our pipeline for two reasons: 1) immune clearance of intracellular pathogens in mouse cells is activated by exogenous IFN-gamma, which we did not add to the host cells, and 2) all known strains of *Toxoplasma* escape IFN-gamma-mediated pathogen clearance by suppressing the IFN-gamma signaling pathway if they infect the host cell before it is exposed to extracellular IFN-gamma (33, 34). In another potential mechanism, an infected host cell may divide and produce an uninfected daughter (a U-Id cell, for <u>U-I</u> by <u>d</u>ivision). To limit the occurrence of U-Id cells, we serum starved all host cells for the 24 hours preceding infection to induce cell cycle arrest (Fig. S3A) and limited infection durations to 3 hpi or less, as live video microscopy of 200 parasite-infected, serum-starved 10 T1/2 host cells for 16 hours revealed that no infected cell divided before 3.67 hpi (Fig. S3B). Finally, a newly internalized parasite might spontaneously exit early from the host cell, a rare and poorly characterized process that is therefore difficult to control. Though we cannot exclude the possibility that a very small number of our U-I cells arose from this mechanism of premature parasite egress, and though U-Id production cannot be completely ruled out, an advantage of scRNA-seq is its ability to distinguish differences between cells arising via different mechanisms (assuming they differ transcriptomically).

### Technical Validation of Single Cell Sequencing for Cells Exposed to or Parasitized by Toxoplasma gondii

Given the limited duration of infection (≤ 3 hours), we expected relatively subtle transcriptional changes in U-I and I-I cells. To maximize the sensitivity of scRNA-seq analyses to detect such signatures, we employed Smart-seq2 library preparation and sequenced to a depth of 1 million reads per cell. All reads were aligned to a concatenated mouse-*Toxoplasma* genome using the genomic sequence for the GT1 parasite strain, a clonal relative of RH (35, 36). In addition, we identified and discarded 43 mouse genes to which reads from extracellular RH parasites aligned (mostly representing evolutionarily conserved genes; Table S1).

To ensure that poorly amplified or poorly sequenced host cells did not confound downstream analysis, we filtered samples based on several quality metrics (Materials and Methods), yielding 453, 2,026, and 2,875 cells at 1, 2, and 3 hpi, respectively, for downstream analysis (Fig. S4A). Mapping efficiency of the analyzed cells was >85% in all experimental trials (Fig. S4B). Characterization of measurement sensitivity based on logistic regression modeling of ERCC spike-in standards revealed a 50% detection rate of 31, 11, and 34 RNA molecules per cell at 1 hpi, 2 hpi, and 3 hpi, respectively (Fig. S4C), amounting to a level of sensitivity comparable to that previously reported (37).

After filtering out genes with expression above the detection limit in <6 cells, we normalized for sequencing depth across cells by dividing each read count by the median read sum to yield counts in units of counts per median (cpm) (Materials and Methods). Gene expression in the scRNA-seq dataset exhibited a strong positive correlation to expression in bulk RNA-seq samples processed the same way specifically for differentially expressed genes (DEGs) between experimental conditions (i.e., U-U, I-I, and U-I cells; Fig. S4D). Furthermore, each bulk experiment’s expression data exhibited the best correlation with their cognate single cell data (Fig. S4E). Taken together, these data demonstrate that scRNA-seq captures similar transcriptomic signatures in host cells to those detected in bulk RNA-seq experiments, validating the scRNA-seq platform as an approach to further characterize host cell transcriptomic responses during *Toxoplasma* infection.

### Single Cell Resolution Reveals Cell-to-Cell Heterogeneity Inaccessible to the Bulk RNA

### Sequencing Platform

A key advantage of single cell resolution is that it enables identification and separation of parasite-independent host cell heterogeneity. Accordingly, single cell resolution facilitates an extra checkpoint to validate the identities of U-I, I-I and U-U cells by quantifying the percentage of *Toxoplasma*-derived reads in each cell. As expected, *Toxoplasma* read content across all cells exhibited a bimodal distribution, where most uninfected cells contained <0.01% *Toxoplasma* reads, while most infected cells possessed higher percentages, i.e., 0.5-4%. However, a small proportion of U-I cells and I-I cells exhibited unexpectedly high (for U-I) or low (for I-I) *Toxoplasma* read counts and were considered to be misclassified (Fig. S5A). This may have resulted from a low rate of TdTomato or mCherry loss in some of the parasites (for cells misclassified as U-I) or from attached but not fully invaded parasites being dislodged from the host cell at some point between fluorescence detection and deposition of the cell into lysis buffer (for cells misclassified as I-I). Such misclassified cells may have contributed significant and potentially misleading signatures to their respective samples. To preclude this possibility in our single cell dataset, we excluded all U-I cells with >0.04% *Toxoplasma* reads and all I-I cells with <0.32% *Toxoplasma* reads from downstream analysis.

Next, because the experimental pipeline examined cells during their earliest interaction with *Toxoplasma*, we expected the subtle, parasite-dependent transcriptional signatures to be potentially eclipsed by intrinsic host processes such as cell cycle, which still progressed in some cells even with serum starvation (Fig. S3A). To separate host cell cycle from other biological processes the parasites could potentially modulate, we determined the phases of all single cells (i.e., G1, S, or G2/M) based on expression of 175 curated cell cycle marker genes (Fig. S5B; Table S2; Materials and Methods). A breakdown of cell cycle phase composition for each experimental condition across all time points revealed a consistent pattern at 2 hpi in which the proportion of cells in G2 or M phase increased in a manner that appeared dependent on injected effectors, i.e., from 13.3% in Mock to 33.9% in Wt U-I, 41.4% in Dmyr1 U-I, and 22.5% in CytD U-I; Fig. S5C). These findings are consistent with potential induction of cell cycle arrest in U-I cells by injected effectors, as previously reported (38–41).

To identify other sources of heterogeneity in the dataset, we used the *Uniform Manifold Approximation and Projection* (UMAP) algorithm for dimensionality reduction and to visualize relationships between all cells based on the most dispersed (i.e., variable) genes. Leiden clustering revealed three distinct clusters, designated herein as populations 1, 2, and 3 (Fig. S5D). Curiously, cells from different time points exhibited differences in the proportion of cells belonging to each population, where 2 hpi cells fell entirely in population 1, and 3 hpi cells fell in all three populations. In conventional single cell analysis, dimensionality reduction and cell clustering are used to identify novel cell types. Though the 10 T1/2 host cell line used in our experiments is clonal, it is derived from a pluripotent stem cell population that has a propensity to differentiate, such that the three identified populations could conceivably represent distinct differentiation states (42).

To limit cell-to-cell heterogeneity from overshadowing potentially subtle transcriptomic signatures induced by parasite effector secretion and parasite invasion, we limited our remaining analyses to host cells in G1 phase (to limit cell cycle signatures) and from UMAP population 1 (to factor out potential cell type signatures).

### Injected Parasite Effectors Drive Inflammatory Transcriptional Signatures Associated with Parasite Infection

Having established a robust pipeline to isolate, RNA sequence, and bioinformatically analyze the transcriptomes of individual cells from each of seven relevant cell species (i.e., Wt U-I, Wt I-I, Wt U-U, Dmyr1 U-I, Dmyr1 I-I, Dmyr1 U-U, and Mock cells; Fig. 1C), we next sought to determine host responses to each of five individual classes of parasite-dependent stimuli, namely rhoptry proteins (ROPs), MYR-dependent dense granule proteins (MDGs), MYR-independent dense granule proteins (MIGs), parasite invasion, and paracrine effects. To isolate each parasite-dependent stimulus, we used the Model-based Analysis of Single Cell Transcriptomics (MAST) algorithm to perform differential gene expression analysis on key pairs of the seven relevant cell species from the pool of G1 phase, UMAP population 1 cells. Across all pairwise comparisons between conditions within each time point, 39, 2,252, and 10,995 differentially expressed genes (DEGs) were detected at 1 hpi, 2 hpi, and 3 hpi, respectively (Table S3A). These results show that parasite effectors and invasion result in almost no detectable transcriptional response at 1 hpi, a modest response involving regulation of a core group of genes at 2 hpi, and ramping up of this response at 3 hpi in which transcription in a much larger set of host cells is modulated. Because of the negligible response at 1 hpi, we excluded data from this time point from the remaining analyses.

For additional validation, we assessed the 2 hpi and 3 hpi datasets for their agreement with two previous bulk RNA-seq studies that captured the host response to infection at 6 hpi in nominally the same parasite strains and culture conditions, albeit in HFFs instead of mouse 10 T1/2 fibroblasts (10, 31). Because these studies measured the infection response by comparing infected vs. mock-infected cells, we examined DEGs between Wt I-I and mock-infected (i.e., Mock) cells in our own datasets. Gene set enrichment analysis (GSEA) of the resulting DEGs at 2 hpi and 3 hpi (Table S3B) using the Molecular Signatures Database’s Hallmark gene sets (43) revealed that nearly all significantly enriched gene sets at 2 hpi and 3 hpi were previously identified in the Naor et al. reference dataset, and many pertained to immune processes (Fig. 3; 31). Of note, gene sets were considered to be significantly enriched if their false discovery rates were < 0.25, a standard cutoff for GSEA given the lack of coherence in most transcriptional datasets and the relatively low number of gene sets being analyzed. Nearly all gene sets *not* preserved in the Naor et al. reference were enriched from genes *down*regulated upon infection. The lack of agreement between our downregulated gene sets and those obtained from the references likely reflects the infection response’s tendency towards gene *up*regulation, and to stochasticity in expression levels for the substantially fewer downregulated genes detected, an interpretation corroborated by the downregulated genes’ higher p-values and lower fold changes. Overall these results indicate that the 2 hpi and 3 hpi datasets capture well-known host responses to infection with *Toxoplasma gondii*.

**Figure 3.**
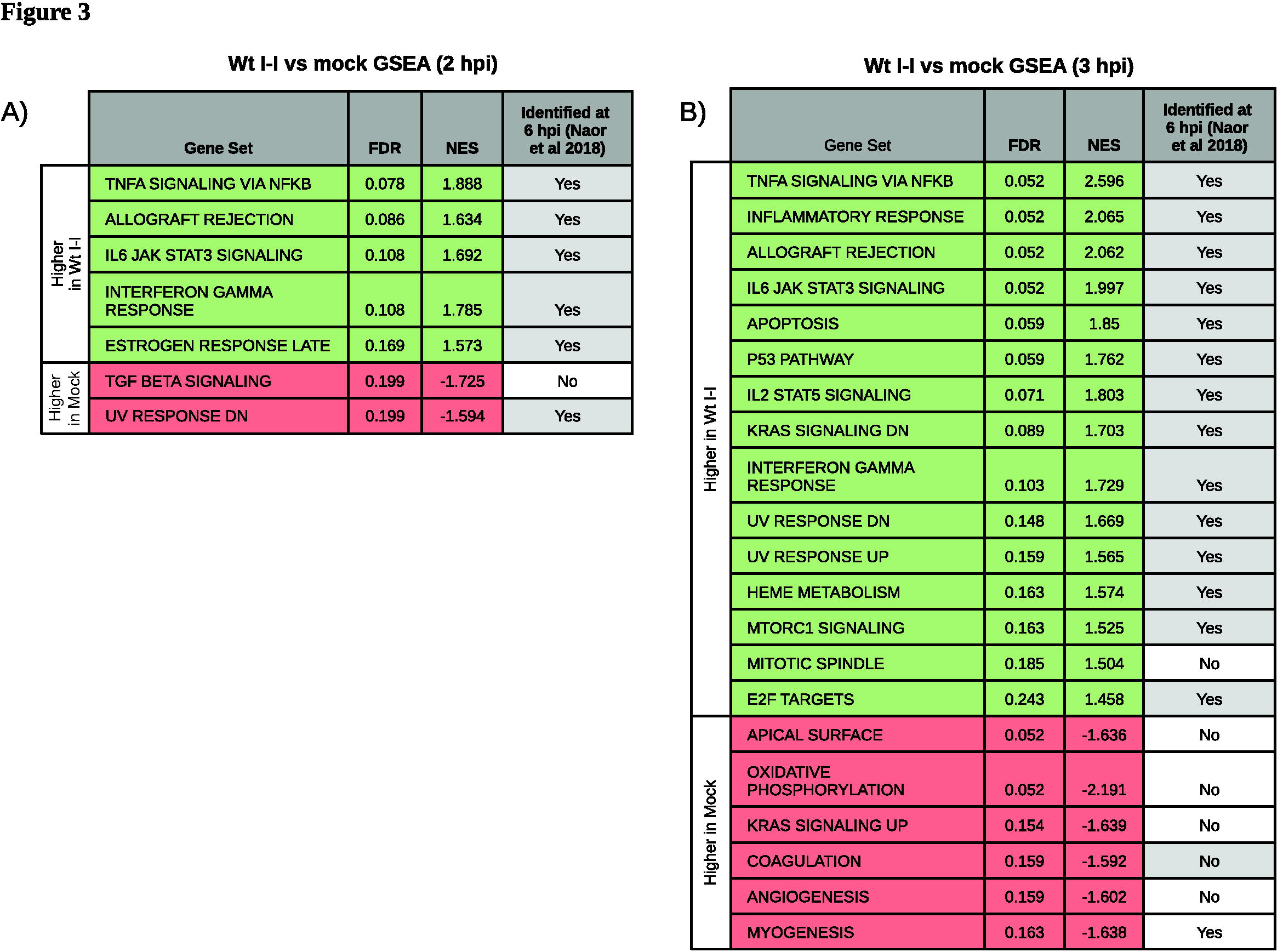
Infection response recapitulates previously identified signatures of early infection. **(A)** Gene set enrichment analysis (GSEA) of the ranked list of differentially expressed genes between Wt I-I and mock cells at 2 hpi (FDR < 0.1, FC > 1.5x) using the Hallmark gene sets from the Molecular Signatures Database. **(B)** GSEA as in (A) of the ranked differentially expressed genes between Wt I-I and mock cells at 3 hpi. FDR = false discovery rate, NES = normalized enrichment score, and FC = fold change. Gray cells indicate pathways common to those identified with FDR <10^-5^ in Naor et al. 2018.

Next, to determine the host response to injection of parasite effectors, we compared U-I cells from wild type parasite infection (i.e., Wt U-I cells) to uninfected cells from the same monolayer (i.e., Wt U-U cells). At 2 hpi, only 10 DEGs were identified (Table S3C), and GSEA of these DEGs revealed no significant enrichment of the Hallmark gene sets. Therefore, injection of parasite effectors without invasion appears to elicit only a trace response at 2 hpi. At 3 hpi, 156 DEGs were detected between Wt U-I and Wt U-U cells (Table S3C), for which GSEA revealed at total of 17 gene sets (Fig. 4A); 14 corresponded to genes expressed higher in Wt U-I cells than in Wt U-U cells, and several were associated with inflammation. In addition, 10 gene sets were common to the infection response, i.e., also enriched in Wt I-I vs. Mock DEGs (Fig. 3), which suggests that much of the early response to parasite infection is driven by the injection of parasite effectors (likely ROPs) into host cells prior to invasion.

**Figure 4.**
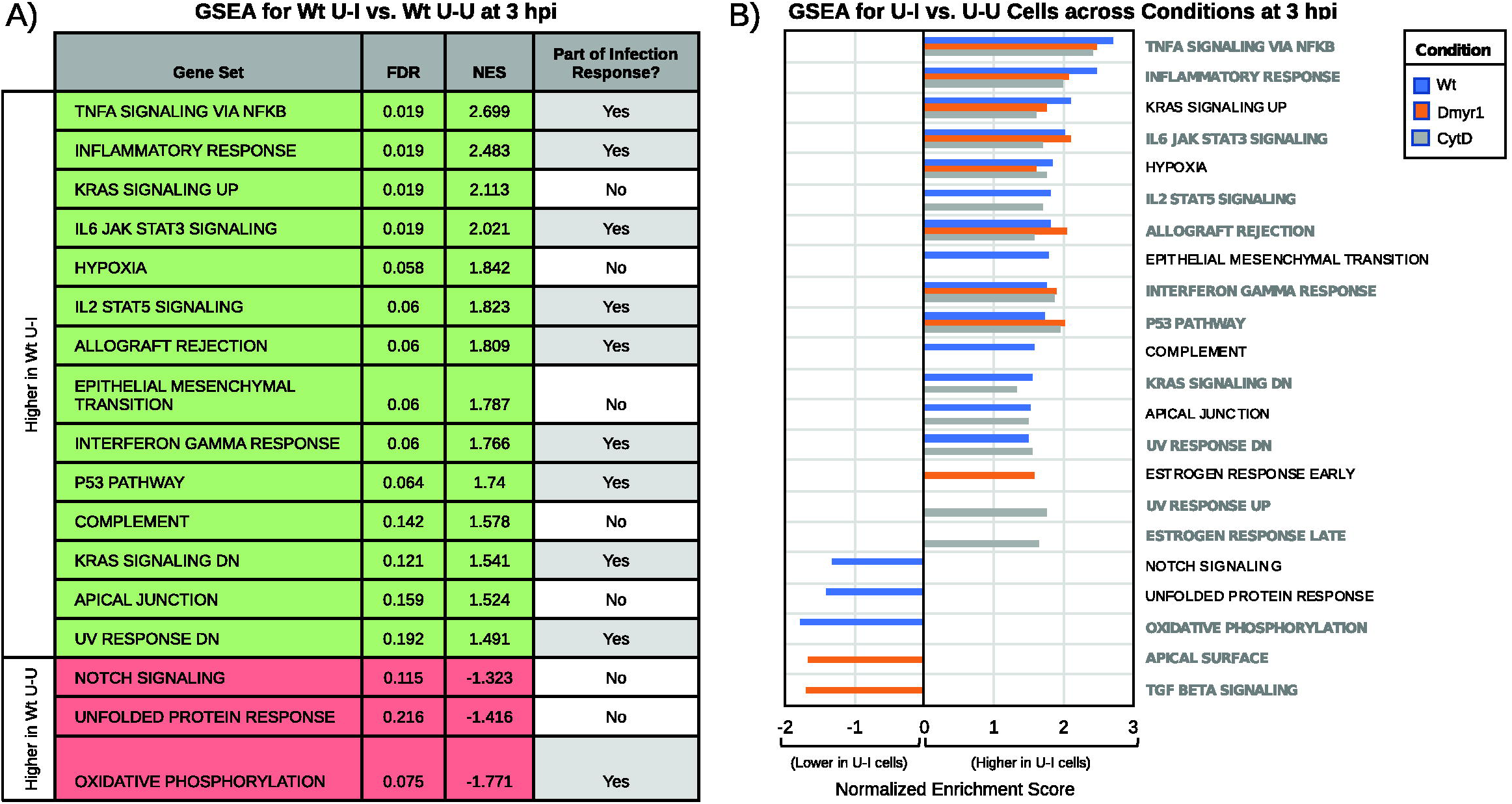
Host response to injected parasite effectors. **(A)** Gene set enrichment analysis (GSEA) of the ranked list of differentially expressed genes (DEGs; false discovery rate (FDR) < 0.1, fold change > 1.5x) between Wt U-I and Wt U-U cells at 3 hpi using the Hallmark gene sets. Green rows indicate gene sets enriched from genes expressed higher in Wt U-I than in Wt U-U, and red rows indicate gene sets enriched in genes expressed lower in Wt U-I than in Wt U-U. Gene sets are designated as part of the infection response if they were also found to be enriched from DEGs between Wt I-I vs. Mock at 2 hpi or 3 hpi. **(B)** GSEA as in (A) between various U-I cells (i.e., from the Wt, CytD, and Dmyr1 infection conditions) and their cognate U-U cells at 3 hpi. Gene sets in bold gray text were also found to be enriched in the infection response. Positive normalized enrichment scores represent enrichment in genes expressed higher in U-I cells than in U-U cells.

To confirm that the transcriptional signatures in Wt U-I cells originated from ROP injection, we made two additional comparisons. For the first, we compared U-I cells from *△myr1* parasite infection (i.e., Dmyr1 U-I cells) to uninfected cells from the same monolayer (i.e., Dmyr1 U-U cells). As MDGs fail to traverse the PVM during *△myr1* parasite infections, the absence of MYR1 should effectively limit the parasites’ effector repertoire to ROPs and MIGs; consequently, transcriptional signatures from Dmyr1 U-I cells and Wt U-I cells should largely resemble one another. In the second comparison, to account for the possibility that Wt U-I signatures originated from a pre-existing difference in those host cells from their neighbors, rather than from injected effectors, we collected U-U, U-I, and I-I cells from host monolayers exposed to wild type parasites pre-treated with the invasion inhibitor cytochalasin D, which allows parasite attachment but blocks subsequent invasion. As expected, cytochalasin D treatment increased the proportion of U-I cells in infected monolayers by ∼6-fold at 2 hpi and by ∼9.3-fold at 3 hpi (Fig. S6), so at least 85-90% of the artificially induced U-I cells (i.e., CytD U-I cells) presumably arose from drug-induced abortion of parasite invasion events, rather than from parasite-independent host cell differences. We predicted that transcriptional signatures detected in CytD U-I cells would be driven almost entirely by injected parasite effectors and would mirror those identified in both Wt U-I and Dmyr1 U-I cells.

As expected, at 2 hpi very few DEGs (29 and 51, respectively) and no enriched gene sets were identified for Dmyr1 and CytD U-I cells vs. their corresponding U-U cells, while at 3 hpi substantially more DEGs (103 and 174 for Dmyr1 and CytD U-I vs. U-U cells, respectively) and gene sets were identified (Table S3D; Table S3E). The 3 hpi gene sets exhibited a strong degree of overlap with the original Wt U-I vs. Wt U-U comparison, such that 14 of the 17 Wt U-I vs. Wt U-U gene sets were also identified in one or both of the corresponding Dmyr1 and CytD comparisons (Fig. 4B). These data are consistent with an inflammatory response to ROP injection that drives much of the host cell’s total response to parasite infection.

In light of the many shared genes and gene sets between the infection vs. injection responses, we next sought to define the distinction between these responses by comparing Wt U-I (injected) to Wt I-I (infected) cells using two complementary approaches. In the first approach, we used MAST to identify DEGs between Wt U-I and Wt I-I cells and subjected the DEGs to GSEA. To better compare the trajectories of Wt I-I and Wt U-I gene expression, we also computed fold changes in DEG expression between each of these conditions and Wt U-U cells (Table S3F). In the second approach, we identified genes with significant differences between Wt U-I and Wt I-I cells in their correlation to a quantity called the CCF2-AM ratio, i.e., the ratio of blue (cleaved) to green (uncleaved) CCF2-AM indicator dye detected during FACS sorting.

In this second approach, the CCF2-AM ratio was used as a quantitative proxy for the influence of parasite-dependent effectors on individual host cells. Briefly, in the strictest sense, the CCF2-AM ratio is a biological readout for penetration of a given host cell by the injected ROP fusion protein Tfn-BLA, as the intracellular CCF2-AM dye in our pipeline is cleaved by the beta-lactamase in Tfn-BLA. Since the host cells should exhibit more or less equal loading of the CCF2-AM substrate, we presumed that the extent of conversion in CCF2-AM signal from green (uncleaved) to blue (cleaved) reflected both the concentration of Tfn-BLA protein introduced into the cell and the amount of time it had spent within the cell. In Wt U-I cells, Tfn-BLA penetration occurs concomitantly with injection of the other ROP effectors; therefore, the CCF2-AM ratio can be interpreted as a quantitative measure for ROP penetration in Wt U-I cells. Wt I-I cells, however, are presumably penetrated not only by ROPs but also by MIGs, MDGs, and the parasites themselves. Accordingly, since the amount of ROPs injected into cells and the time ROPs spend inside cells likely track with the same quantities for the remaining parasite-dependent stimuli, we used the CCF2-AM ratio in Wt I-I cells as a proxy for the presence of all four parasite stimuli inside each cell. Next, we computed the Spearman correlation between each gene’s expression and the CCF2-AM ratio separately in both Wt U-I cells and Wt I-I cells. Of note, because we calculated the correlation by incorporating the CCF2-AM ratios from Wt U-U cells (which should be devoid of ROPs, MIGs, MDGs, and parasites) as negative controls for both the Wt U-I and Wt I-I analyses, the correlation data do not encapsulate the influence of parasite-dependent stimuli that Wt U-U cells have in common with Wt U-I and Wt I-I cells, i.e., paracrine factors. Finally, we identified genes that exhibited significant shifts between Wt U-I and Wt I-I cells in their relationship to the CCF2-AM ratio (Table S4A) by first modeling a Gaussian distribution of their Spearman correlations to the CCF2-AM ratio (Fig. 5A).

**Figure 5.**
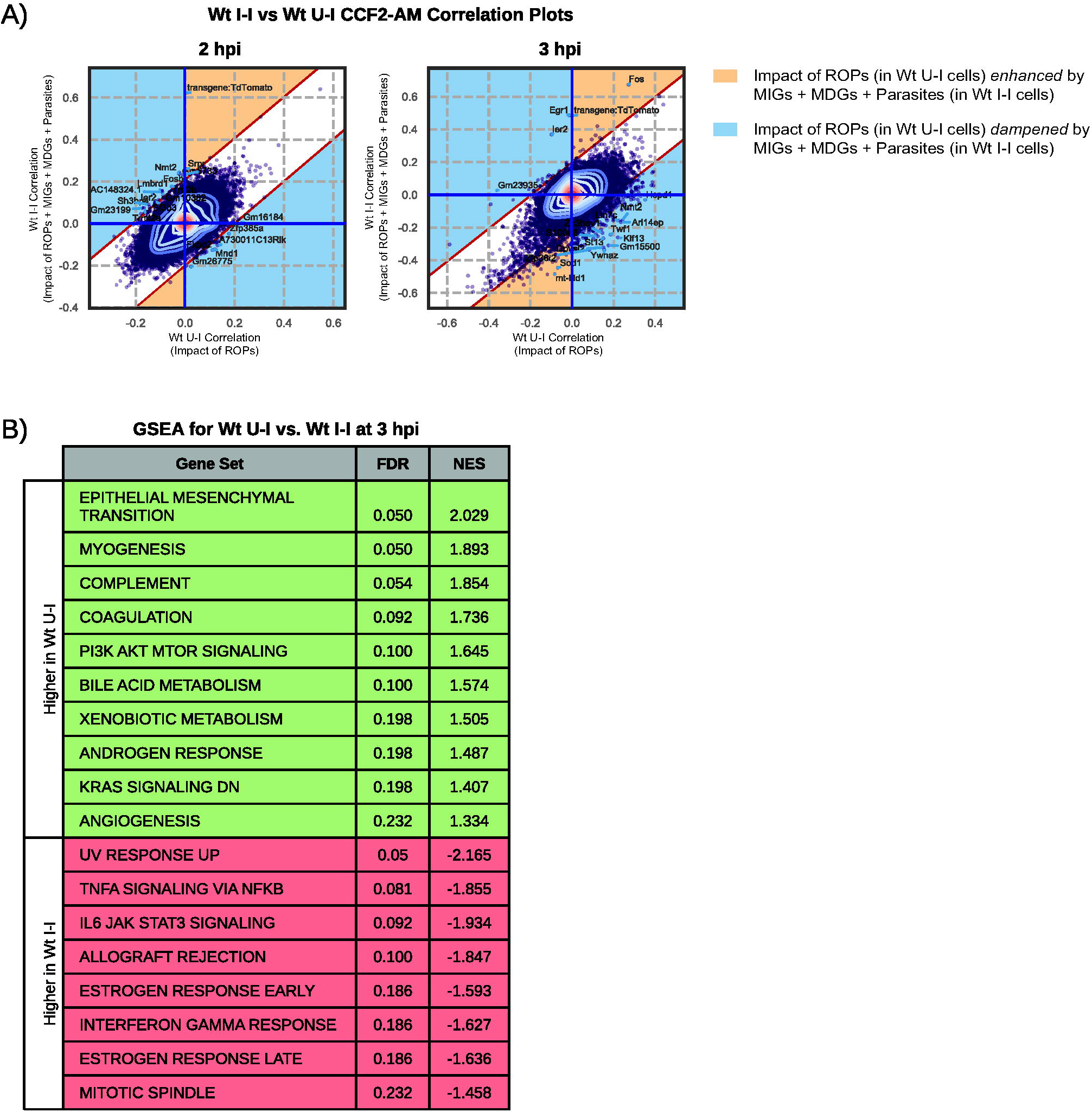
Comparison of host response to injection vs. infection (i.e., signatures in Wt U-I vs. Wt I-I cells.) **(A)** Scatterplot comparing injection-associated genes in Wt I-I cells and Wt U-I cells, as determined by the Spearman correlation of gene expression to the CCF2-AM ratio in Wt I-I and Wt U-I cells. Contours reflect density of points, with central contours being in the area of highest density. Labeled points on the plot are the top 20 genes with the lowest probability of belonging to a Gaussian model fit with parameters that best describe the observed Spearman correlation data. Yellow and blue shading mark regions of the scatterplots where gene expression trends in Wt U-I cells are either enhanced or dampened upon parasite invasion and release of dense granule effectors into the host cell (as seen in Wt I-I cells). **(B)** GSEA of the ranked list of differentially expressed genes between Wt U-I and Wt I-I cells at 3 hpi, where green rows denote gene sets enriched from genes expressed higher in Wt U-I cells than in Wt I-I cells, and red rows represent genes expressed higher in Wt I-I than in Wt U-I. FDR = false discovery rate, NES = normalized enrichment score.

Genes that significantly deviated from the Gaussian distribution and that also exhibited a sufficient difference in their CCF2-AM correlation scores in Wt U-I vs. Wt I-I cells were interpreted to be associated with either ROP injection alone (in Wt U-I cells) or with all secreted effectors plus parasites (in Wt I-I cells, in which ROP penetration should be accompanied by all other parasite-induced insults).

Analysis of Wt I-I vs. Wt U-I cells at 2 hpi revealed no significant DEGs, pointing to a profound similarity in host transcription in Wt I-I and Wt U-I cells at 2 hpi. However, the corresponding CCF2-AM correlation data exhibited a higher sensitivity for detecting differences between Wt I-I vs. Wt U-I cells and yielded 81 significant deviants from the Gaussian distribution (Fig. 5A, left panel). Of the deviants, 73 (∼90.1%) fell in regions of the scatterplot where genes were either: 1) negatively correlated with the CCF2-AM ratio in one condition and positively correlated in the other; or 2) had stronger positive or negative correlations in Wt U-I cells than in Wt I-I cells as determined by a Gaussian model (Fig. 5A, left panel, blue regions; Table S4A). This provides evidence for downstream events in infection (i.e., release of dense granule proteins, parasite invasion) suppressing effects of genes induced by ROP injection as early as 2 hpi.

At 3 hpi, 122 DEGs and 423 CCF2-AM correlation deviants were identified for Wt I-I vs. Wt U-I cells, pointing to a divergence in the injection vs. infection responses at this later time point. Of the DEGs, 100 (∼82.0%) exhibited opposing trajectories in Wt I-I vs. Wt U-I cells (Table S3F), i.e., they were either: 1) upregulated in one cell type and downregulated in the other compared to a common Wt U-U standard; or 2) more upregulated or downregulated in Wt U-I cells compared to the standard than in Wt I-I cells. In addition, of the CCF2-AM correlation deviants, 333 (∼79%) exhibited at least a moderate negative correlation with the CCF2-AM ratio in Wt I-I cells, an observation consistent with these genes being downregulated in response to the combination of all parasite-derived insults and effectors during infection (Fig. 5A, right panel; Table S4A). Furthermore, 282 (∼67%) of the deviants fell in regions of the scatterplot in which the impact of ROPs is dampened by dense granule proteins (GRAs) and parasite invasion (Fig. 5A, right panel, blue regions; Table S4A), which is consistent with host responses in Wt U-I cells being counteracted by parasite effectors that are operating in Wt I-I cells only. This lends further support to the notion that effectors injected circa invasion (i.e., ROPs) are counterbalanced by subsequently introduced effectors (i.e., GRAs). Of note, the remaining 141 CCF2-AM deviant genes at 3 hpi fell within the sections of the scatterplot in which the effects of ROPs are enhanced by those of the GRAs and parasite invasion (Fig. 5A, right panel, yellow regions; Table S4A). This suggests that while GRA release and parasite penetration may counterbalance some ROP-induced host responses, these events may also enhance other genes induced by ROP injection.

To discern the biological significance of the differences between the infection vs. injection responses, we performed GSEA on Wt U-I vs. Wt I-I DEGs. The resulting enriched gene sets (Fig. 5B) included many that were previously identified in the host infection response, with gene sets enriched from genes expressed higher in Wt I-I cells corresponding to more inflammatory processes. Because the vast majority of genes with higher expression in Wt U-I cells than in Wt I-I cells exhibited evidence of ROP effectors being counteracted by subsequently secreted effectors such as MDGs and MIGs (84 out of 93 genes; Table S3F), these findings are consistent with injection-associated inflammatory host processes ramping up upon parasite penetration (and GRA release), while other injection-associated processes are dampened by these later events.

### Parasite Effectors Counteract One Another to Yield a Modest Host Response to Early Parasite Infection

Next, we sought to determine the contribution of individual parasite effector compartments to the difference between the host infection (in Wt I-I cells) vs. ROP injection (in Wt U-I cells) responses. Because Wt U-I and Wt I-I cells are both injected with ROPs and originate from the same monolayer, ROPs and paracrine effectors likely do not explain the differences between these two cell types. Instead, MDGs, MIGs, and the act of parasite penetration itself are, *a priori,* most likely to explain these differences. Our dataset, which includes infected (I-I), bystander uninfected (U-U), and U-I cells originating from monolayers infected with wild type and *Δmyr1* parasites, presents a unique opportunity to examine the impacts of these compartments in isolation.

To determine how individual effector compartments contribute to the transcriptomic differences between Wt I-I vs. Wt U-I cells, we made two key comparisons. In the first comparison, we examined Dmyr1 U-I cells, which presumably respond to ROP injection, and Dmyr1 I-I cells, which are thought to respond to ROP injection plus MIGs and parasite penetration (Fig. 1C; Fig. 1D). We predicted that MIGs and parasite penetration would account for the differences between Dmyr1 I-I and Dmyr1 U-I cells, and for a subset of the differences between Wt U-I and Wt I-I cells. In the second comparison, we examined Wt I-I vs. Dmyr1 I-I cells, which originate from monolayers infected with wild type and *Δmyr1* parasites, respectively. Signatures detected in Dmyr1 I-I cells likely reflect the host response to the combination of ROP injection, MIG activity, parasite penetration of host cells, and paracrine effectors secreted into the extracellular milieu, while those detected in Wt I-I cells likely reflect the response to these elements plus MDG secretion (Fig. 1C). Therefore, we predicted that comparing Dmyr1 I-I vs. Wt I-I would illustrate the impact of MDGs, as well as any paracrine effects dependent on the presence of MYR1 (Fig. 1D), and that this would account for a second subset of the differences between the infection vs. injection response showcased in the Wt I-I vs. Wt U-I comparison.

### MYR-Independent GRAs and Parasite Penetration Collectively Enhance ROP-induced

### Inflammatory Responses

To examine the host response to the combination of MIGs and parasite penetration, we compared Dmyr1 U-I cells, which should be penetrated by ROPs, and Dmyr1 I-I cells, into which parasites should secrete ROPs and MIGs, but not MDGs (Fig. 1C; Table S3G). At 2 hpi, only 5 DEGs and no GSEA enriched gene sets were identified, implying profound similarity between Dmyr1 U-I and Dmyr1 I-I cells at this time point (as was also the case for Wt U-I vs. Wt I-I cells). At 3 hpi, 80 DEGs were identified, pointing to a slight divergence between these two cell types at this time point. 48 (60%) of the DEGs exhibited evidence of being influenced by effectors that counteract one another’s effects (i.e., their expression exhibited either: 1) opposing trends in Dmyr1 U-I vs.

Dmyr1 I-I cells when compared to an uninfected Dmyr1 U-U standard; or 2) a stronger induction or suppression in Dmyr1 U-I cells than in Dmyr1 I-I cells), which suggests that MIGs may play a role in neutralizing the effects of the ROPs preceding them. Note, however, that we cannot exclude the possibility that this effect is attributable to stimuli related to physical penetration by the parasites.

GSEA of the 3 hpi DEGs identified 15 enriched gene sets (Fig. 6), of which 9 were also found to be enriched in the Wt U-I vs. Wt I-I comparison (Fig. 5B). For gene sets enriched from genes expressed higher in Dmyr1 I-I cells than in Dmyr1 U-I cells, nearly all corresponded to gene sets already identified as part of the host response to ROP injection and/or were associated with inflammatory processes. In contrast, none of the gene sets enriched from genes expressed higher in Dmyr1 U-I cells were identified as part of the injection response and instead corresponded to other processes, i.e., complement cascade, coagulation, MTORC1 signaling, and myogenesis. Taken together, these results suggest that MIGs and parasite penetration do indeed account for some of the difference between the infection response (in Wt I-I cells) vs. the injection response (in Wt U-I cells). More specifically, genes corresponding to inflammatory processes that are already induced upon ROP injection (in Wt U-I and Dmyr1 U-I cells) appear to be further induced, i.e., enhanced, by the combination of MIGs and parasite penetration, while genes for which the effect of MIGs + Parasites dampen the influence of the ROPs appear to correspond to a different set of cellular processes. Although this latter set of genes accounts for the majority (60%) of Dmyr1 U-I vs. Dmyr1 I-I DEGs, the majority of the total enriched gene sets corresponds to cellular processes enhanced rather than dampened by MIGs + Parasites, and these enhanced gene sets collectively exhibit much higher statistical significance. This raises the possibility that at least some processes dampened by MIGs + Parasites may not be adequately captured by the Hallmark gene sets.

**Figure 6.**
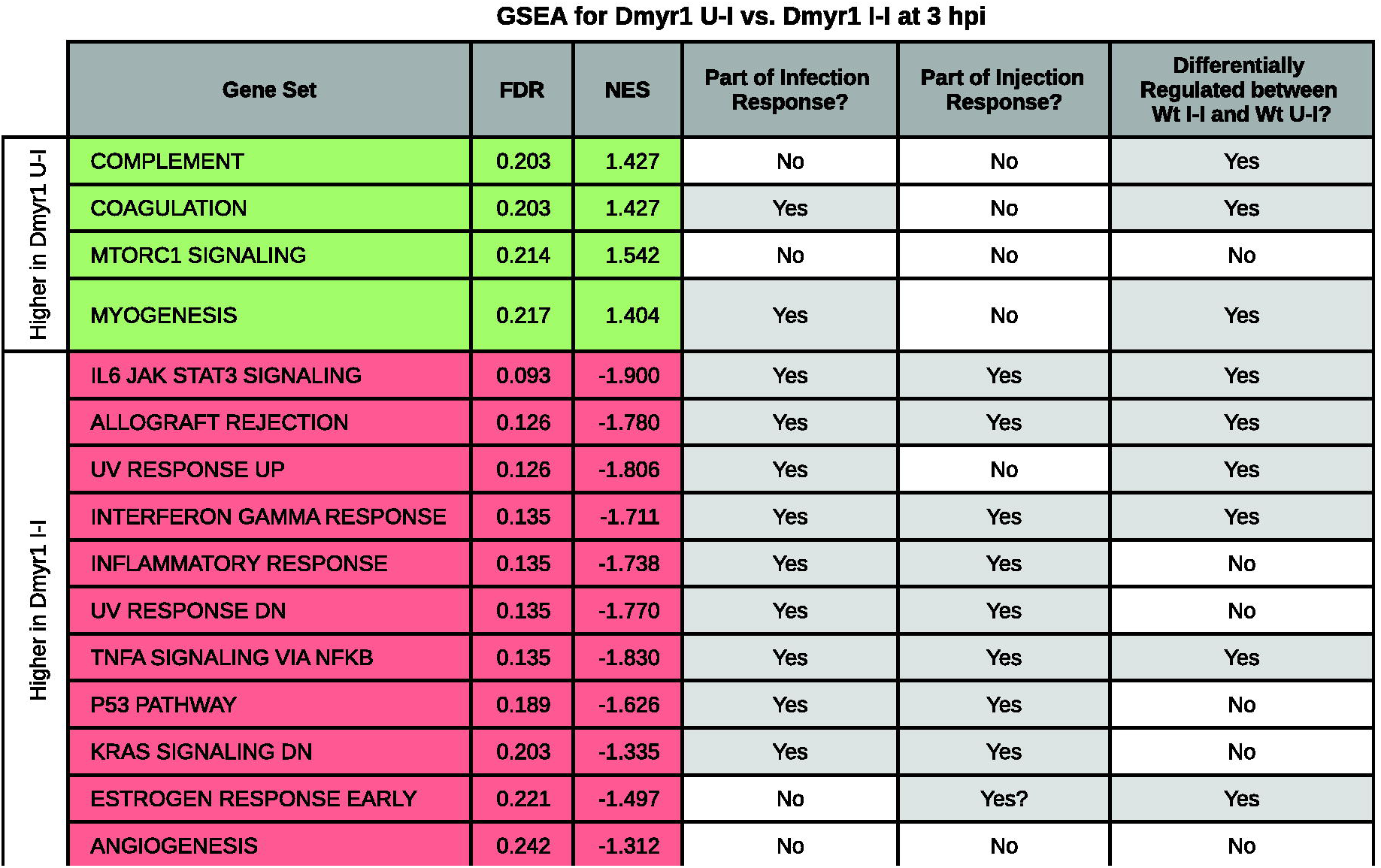
GSEA of the ranked list of differentially expressed genes between Dmyr1 U-I and Dmyr1 I-I cells at 3 hpi, which illustrates the impact of MYR-independent dense granule proteins (MIGs) and parasite invasion on the host response to infection with *Toxoplasma.* Green rows denote gene sets enriched from genes expressed higher in Dmyr1 U-I cells than in Dmyr1 I-I cells, and red rows represent genes expressed higher in Dmyr1 I-I cells than in Dmyr1 U-I cells. FDR = false discovery rate, NES = normalized enrichment score.

### MYR-Dependent GRAs Counterbalance Parasite Effectors Released Earlier in the Lytic

### Cycle

A previously published bulk RNA-seq experiment comparing host transcription in monolayers infected with wild type vs. *Δmyr1* parasites at 6 hpi (31) revealed a set of genes such that in the absence of MYR1, expression changes were unmasked, while in the presence of MYR1 (i.e., during wild type infection), there was no net change. This work implies that collectively, MDGs and associated paracrine effectors secreted during infection with wild type but not *Δmyr1* parasites participate in counterbalancing transcriptional signatures induced by prior parasite-dependent stimuli.

To determine whether MDGs and associated paracrine effects play a similar role in the present single cell datasets, and to ascertain their contribution to the difference between the infection vs. injection responses, we compared Dmyr1 I-I and Wt I-I cells. MAST analysis of this comparison identified 46 and 367 DEGs at 2 hpi and 3 hpi, respectively (Table S3H). While GSEA of the 2 hpi DEGs returned no significantly enriched gene sets, GSEA at 3 hpi revealed 16 significantly enriched gene sets, of which all but one were enriched from genes expressed higher in Dmyr1 I-I cells than in Wt I-I cells (Fig. 7A). Furthermore, 8 gene sets were identified in the previously published bulk RNA-seq experiment comparing infections with wild type vs. *△myr1* parasites (31), while 6 corresponded to infection-associated gene sets (i.e., were enriched from Wt I-I vs. Mock DEGs), and 8 were identified as part of the ROP injection response (i.e., were enriched from Wt U-I vs. Wt U-U DEGs). Taken together, these data are consistent with MDGs and/or their associated paracrine effects selectively impinging on Wt I-I cells and dampening the effects of parasite effectors introduced into host cells earlier in the lytic cycle. The overlap between these gene sets and those of the injection response shows that some of the effectors whose responses were dampened by MDGs and associated paracrine effects include injected effectors (i.e., ROPs), which was previously suspected (31) but never before explicitly demonstrated.

**Figure 7.**
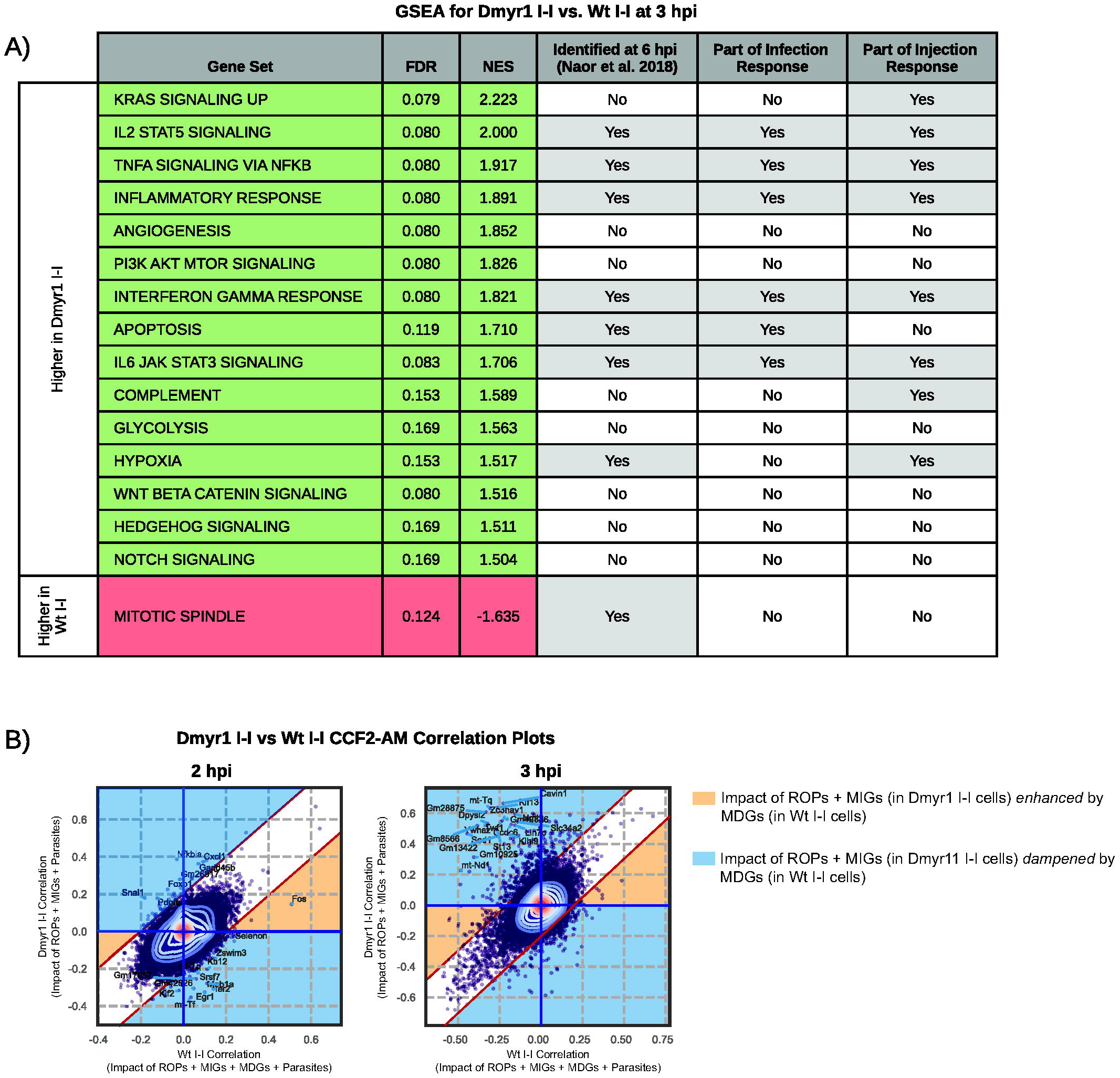
Impact of MYR-dependent dense granule proteins (MDGs) and MYR-dependent paracrine effectors, as illustrated by comparing Dmyr1 I-I and Wt I-I cells. **(A)** Gene set enrichment analysis (GSEA) of ranked list of differentially expressed genes between Dmyr1 I-I and Wt I-I cells at 3 hpi. Green rows denote enrichment from genes expressed higher in Dmyr1 I-I cells than in Wt I-I cells, and red rows represent enrichment from genes expressed higher in Wt I-I than in Wt U-I. FDR = false discovery rate, NES = normalized enrichment score. **(B)** Scatterplot comparing injection-associated genes in Dmyr1 I-I cells and Wt I-I cells, as determined by the Spearman correlation of gene expression to the CCF2-AM ratio in Dmyr1 I-I and Wt I-I cells. Contours reflect density of points, with central contours being in the area of highest density. Labeled points on the plot are the top 20 genes with the lowest probability of belonging to a Gaussian model fit with parameters that best describe the observed Spearman correlation data. Yellow and blue shading mark regions of the scatterplots where gene expression trends in Dmyr1 I-I cells are either enhanced or dampened by MDGs or paracrine effects (as seen in Wt I-I cells).

Though comparing Dmyr1 I-I vs. Wt I-I cells illustrates the collective impact of MDGs and MYR-dependent paracrine effects on host transcription (Fig. 1D), this comparison cannot distinguish the individual impacts of these two stimuli. To determine the effect of MDGs alone, we performed CCF2-AM ratio correlation analysis on Wt I-I vs. Dmyr1 I-I cells (Table S4B), as was performed for Wt I-I vs. Wt U-I. As was the case for computation of both Wt I-I and Wt U-I CCF2-AM correlations, this type of analysis excludes the impact of paracrine effects because each set of CCF2-AM correlation data is computed using the CCF2-AM ratios of both I-I cells and U-U cells from the same infected monolayer. 224 and 343 CCF2-AM ratio correlation deviants were identified between Wt I-I and Dmyr1 I-I cells at 2 hpi and 3 hpi, respectively. 218 (∼97.3%) and 297 (∼86.6%) deviants at 2 hpi and 3 hpi, respectively, fell in the regions of the CCF2-AM correlation scatterplot in which the impact of effectors found in Dmyr1 I-I cells (i.e., ROPs + MIGs) are dampened by effectors found in Wt I-I cells (i.e., MDGs; Fig. 7B, blue regions; Table S4B), which implies that signatures induced by effectors released into Dmyr1 I-I cells are counteracted specifically by MYR-dependent effectors released into Wt I-I cells.

### MYR-Dependent Paracrine Effects Also Counterbalance Parasite Effectors Released

#### Earlier in the Lytic Cycle

To determine the impact of specifically paracrine factors on the host response, we examined two key comparisons: Wt U-U vs. Mock (which captures all paracrine effects; Table S3I) and Dmyr1 U-U vs. Mock (which captures MYR-independent paracrine effects; Table S3J). As illustrated in Fig. 1C and Fig. 1D, differences between these two comparisons should be attributable to paracrine factors released from host cells in a MYR-dependent fashion.

At both 2 hpi and 3 hpi, paracrine effects in *Δmyr1* parasite infections exhibited robust differences from paracrine effects in wild type infections. At 2 hpi, ∼196 and 54 DEGs were identified for the Dmyr1 and Wt cell comparisons, respectively; at 3 hpi, the difference was even more pronounced, with 1,864 DEGs identified for the Dmyr1 comparison, vs. only 20 DEGs for Wt. In addition, GSEA of Dmyr1 U-U vs. Mock DEGs exposed an abundance of gene sets, many of which corresponded to inflammatory processes and other pathways found to be part of the infection response, whereas GSEA of Wt U-U vs. Mock DEGs enriched for few, if any gene sets (3 and 0 at 2 hpi and 3 hpi, respectively; Fig. 8A). Furthermore, the responses of these DEGs were reproduced consistently between not only U-U cells vs. mock-infected cells, but also between U-I or I-I cells vs. mock-infected cells (Fig. 8B), which establishes that these trends affect all cell types within a given infected monolayer. Taken together, these results suggest that gene expression trends induced during *Toxoplasma* infection (i.e., those encapsulated by the DEGs that arise from comparing Dmyr1 U-U and Mock cells), including those corresponding to pathways that respond to injected ROPs, are suppressed via a MYR-dependent paracrine mechanism. Accordingly, supernatants taken from host cell cultures infected with wild type and *Δmyr1* could, in theory, exert transcriptional influence on fresh host cell monolayers, an interesting avenue for future investigation.

**Figure 8.**
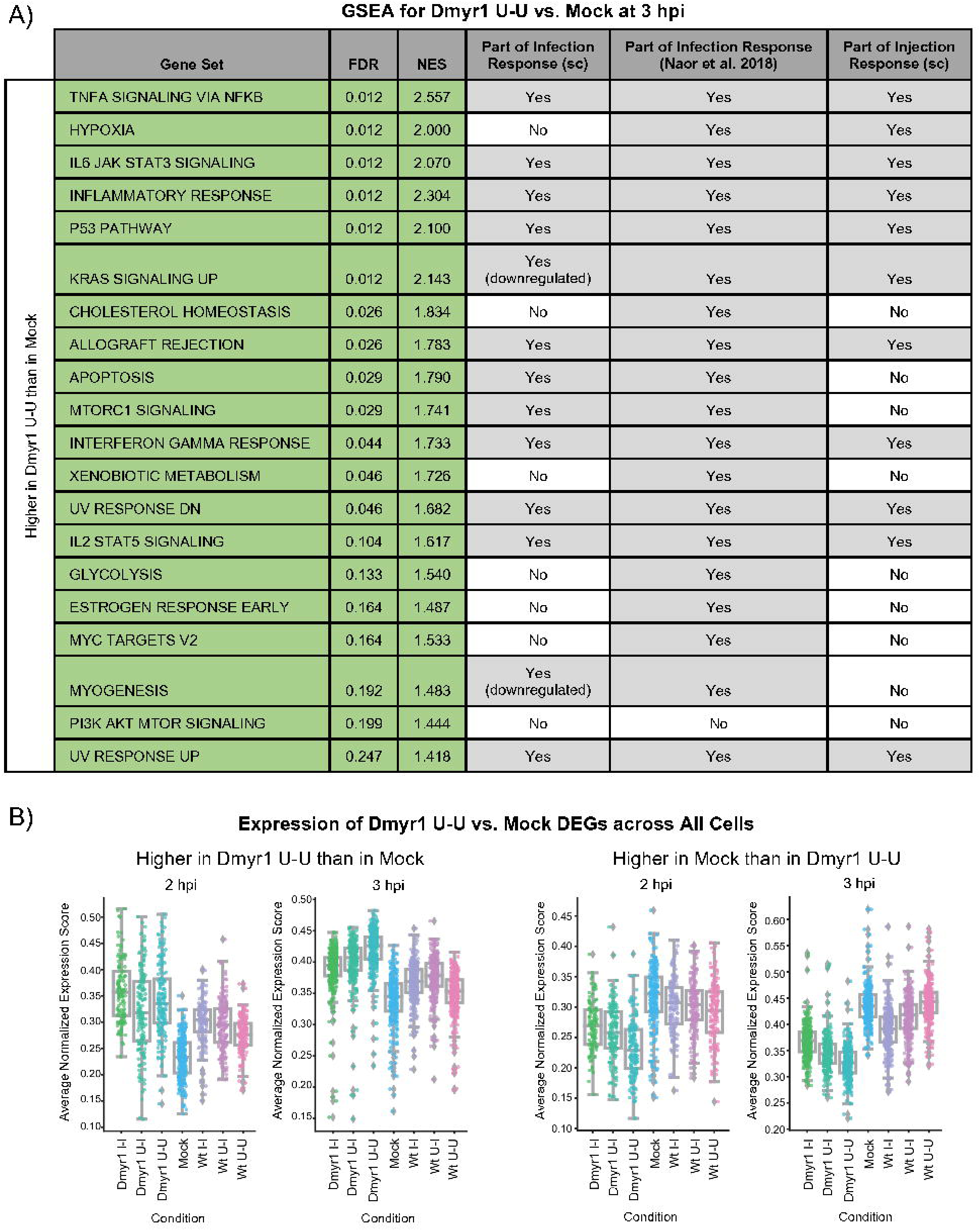
Comparison of Dmyr1 U-U vs. Mock cell types reveals impact of MYR-dependent paracrine effects on host cell transcription during *Toxoplasma* infection. **(A)** Gene set enrichment analysis (GSEA) of the ranked list of differentially expressed genes between Dmyr1 U-U and Mock cells at 3 hpi. FDR = false discovery rate, NES = normalized enrichment score, and sc = the current single cell RNA-seq dataset. **(B)** Trends in expression of Dmyr1 U-U vs. Mock DEGs are conserved in U-I and I-I cells from Wt and Dmyr1 infections, as illustrated by strip plots depicting expression of DEGs between Dmyr1 U-U and Mock cells across all cell types. Each point represents a single cell, and its y-axis displacement reflects the average of the normalized expression scores for all DEGs between Dmyr1 U-U and Mock cells. The normalized expression score of a given gene is calculated by scaling the log_2_ count per median (cpm) for that gene, such that the cell with the lowest cpm receives a score of 0, and the cell with the highest cpm receives a score of 1.

### Model of Host Responses to Individual *Toxoplasma* Parasite-Dependent Stimuli and Effector Compartments

The preceding analyses have accounted for host responses to 5 parasite-dependent stimuli: 1) rhoptry protein (ROP) injection; 2) MYR-independent dense granule (MIG) secretion; 3) MYR-dependent dense granule (MDG) secretion; 4) paracrine effects (which can be further subdivided into MYR-independent vs. MYR-dependent paracrine effects); and 5) parasite invasion. To succinctly represent the interplay between these stimuli, we curated a list of gene sets that best represented the expression trends captured in our analyses. Gene sets were included if they were significantly enriched (false discovery rate < 0.25) from DEGs between a majority of the following 8 key comparisons: 1) Wt U-I vs. Wt U-U (ROP injection); 2) Wt I-I vs. Dmyr1 I-I (MDGs + MYR-dependent paracrine effects); 3) Wt U-U vs. Dmyr1 U-U (MYR-dependent paracrine effects); 4) Dmyr1 I-I vs. Dmyr1 U-I (MIGs + Parasite invasion); 5) Dmyr1 U-U vs. Mock (MYR-independent paracrine effects); 6) Wt I-I vs. Wt U-I (MIGs + MDGs + parasite invasion); 7) Wt I-I vs. Wt U-U (all stimuli except paracrine effects); and 8) Wt I-I vs. Mock (all stimuli). The 12 gene sets selected included those pertaining to immune responses, cell proliferation, cellular stress, and the complement pathway.

Expression trends for the DEGs within these gene sets are summarized in Fig. 9. Based on our analyses, we propose the following model of parasite-driven effects on host cell transcription. First, because Wt U-I cells appeared to induce DEGs in nearly all the showcased gene sets compared to Wt U-U cells, ROP injection likely induces many of the pathways these gene sets capture, particularly immune-related and cellular stress pathways (Fig. 9A and Fig. 9B, ROPs; Fig. 9C, ROP injection). These signatures may arise in response to the ROPs themselves, or due to cell trauma secondary to perforation of the host cell membrane by the parasite during ROP injection. Next, parasites penetrate the host cell and secrete MIGs into the PVM. The host response to these two stimuli together is relatively mild, as evidenced by the high false discovery rates and small number of genes per gene set, even at 3 hpi. Though MIGs + Parasites appeared to counteract the effects of ROP injection at the *gene* level, their most significant impact at the *gene set* level appears to be enhancement of inflammatory transcriptional signatures induced by ROP injection (Fig. 9A and Fig. 9B, MIGs + Invasion; Fig. 9C, MIG Secretion and Parasite Penetration). This does not exclude the possibility that other host processes not covered by the Hallmark gene sets are suppressed by MIGs + Parasites. Next, in cells infected with *Δmyr1* parasites, MYR-independent paracrine factors secreted by neighboring infected cells appear to enhance the effects of ROP injection even more so than MIGs + Parasites and may do so not only for immune-related genes, but also for genes pertaining to cellular stress, complement, and cellular proliferation (Fig. 9A and Fig. 9B, dmyr1UU-mock/P(MI)). In contrast, during wild type infections, two additional parasite-dependent stimuli, MDGs and MYR-dependent paracrine factors, both appear to rein in transcriptional signatures induced by the other stimuli (Fig. 9A and Fig. 9B, MDGs + P(MD); Fig. 9C, MYR-dependent stimuli; Fig. 7B). Together, all five parasite classes produce transcriptional signatures that veer toward induction of genes pertaining to inflammation and cellular stress but that are less pronounced than the response to injection of ROPs.

**Figure 9.**
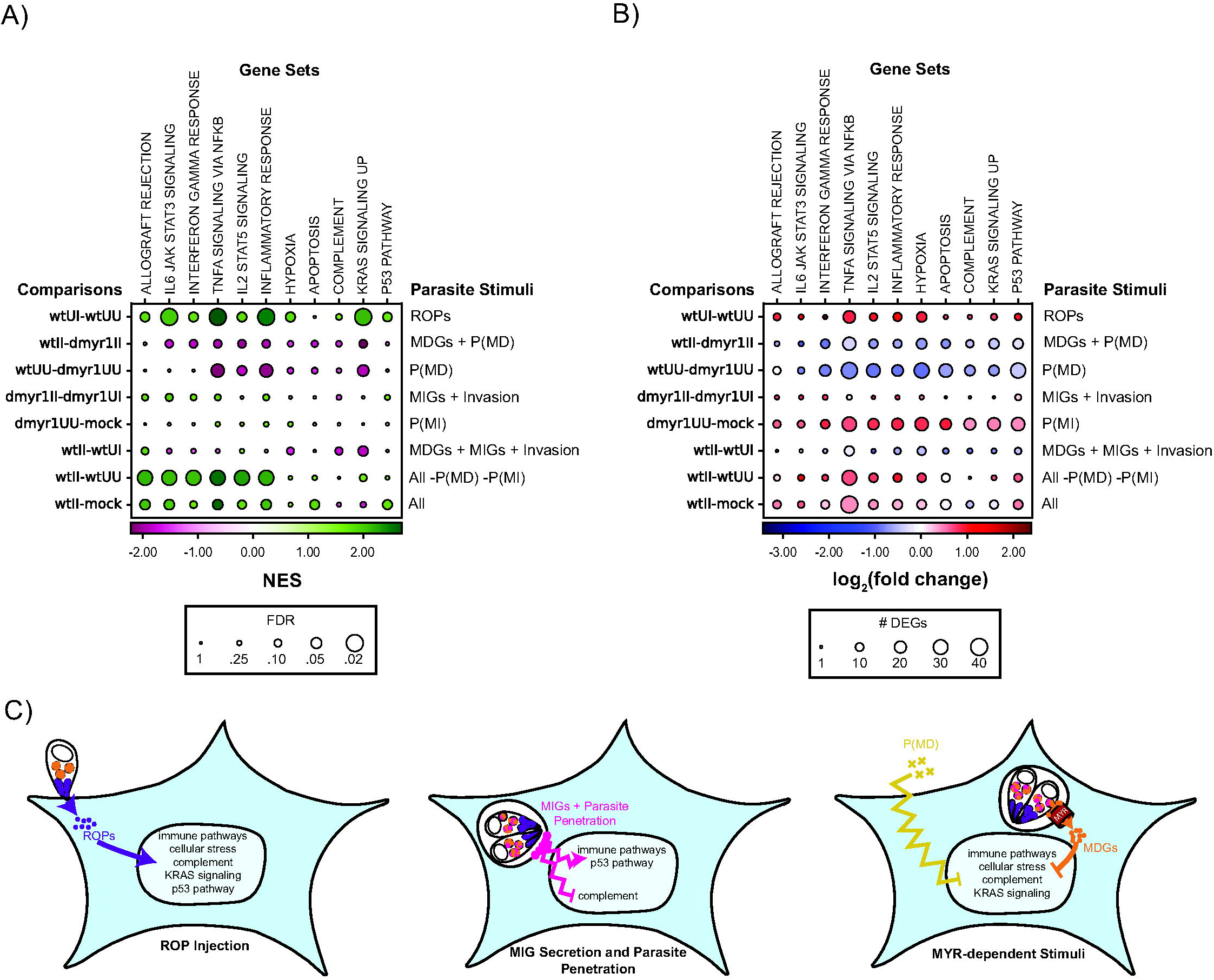
Summary of host transcriptional response to parasite-dependent stimuli. **(A)** Bubble plot of parasite-dependent host transcriptional signatures at 3 hpi that depicts the behavior of 11 representative Hallmark gene sets from the Molecular Signatures Database (columns) across 8 key comparisons between experimental conditions (rows, indicated on left of each plot) that each encapsulates the impact of certain parasite-dependent stimuli on the 11 gene sets (indicated on right of each plot). Colors indicate normalized enrichment scores (NES) from gene set enrichment analysis (GSEA) of differentially expressed genes (DEGs) between each comparison. FDR = false discovery rate, where FDR < 0.25 is considered significant (as is standard for GSEA). Parasite-dependent stimuli include ROPs = rhoptry proteins, MDGs = MYR-dependent dense granule proteins, MIGs = MYR-independent dense granule proteins, P(MD) = MYR-dependent paracrine effects, P(MI) = MYR-independent paracrine effects, and All = ROPs + MDGs + MIGs + Invasion + P(MD) + P(MI). **(B)** Bubble plot as in (A) where colors indicate log_2_ normalized fold change in expression of DEGs that fall under each comparison and each gene set. **(C)** Model of individual impacts of parasite-dependent stimuli on host transcription, which include immune pathways (e.g., TNFA Signaling via NFKB, IL6 JAK STAT3 Signaling, Inflammatory Response, IL2 STAT5 Signaling, Interferon Gamma Response, and Allograft Rejection), cellular stress (e.g., Hypoxia, Apoptosis), and cell proliferation-associated pathways (i.e., KRAS signaling, p53 pathway). Curved lines indicate modulation of host transcription, and jagged lines indicate modulation without effector translocation to the nucleus.

## Discussion

In this study, we examined host responses to infection with the parasite *Toxoplasma gondii* using scRNA-seq on *in vitro* infected, uninfected, and uninfected-injected (U-I) host cells, the latter of which arise from aborted invasion events and that have until very recently (44) been characterized primarily morphologically. Key fixtures of our experimental pipeline included: 1) early time points, to limit isolating false positive U-I cells arising from mechanisms besides aborted invasion; 2) FACS sorts, which purified rare U-I cells and relatively rare infected cells at early time points; and 3) single cell resolution, which enabled bioinformatic validation of all cells’ infection status. The level of confidence lent by these measures to the validity of the captured U-I cells enabled interrogation of host responses specifically to ROP injection, an aspect of parasite infection previously inaccessible to study due to the rapid kinetics of effector secretion at the time of invasion. Note that while others have also used scRNA-seq to measure host responses to *Toxoplasma* infection at a single cell level, these studies analyzed cells from animals at many days post-infection, and therefore did not assess the earliest impacts of infection or particularly ROP injection (45).

Because the experimental pipeline also leveraged infections with *△myr1* parasites, our dataset is a comprehensive resource for the individual impacts of not only ROPs, but also MIGs, MDGs, and paracrine stimuli on host transcription. Our analyses revealed an early response to *Toxoplasma* infection with subtle yet clear signatures overlapping with inflammatory and cellular stress signaling axes. Induction of these axes appears to be 1) driven primarily by ROP injection; 2) enhanced somewhat by the combination of MIGs, parasites, and MYR-independent paracrine factors; and 3) counterbalanced by MDGs and downstream paracrine effects, i.e., factors secreted during wild type but not *Δmyr1* infection. These findings substantiate previous evidence that at least some MDGs suppress host responses induced by other parasite-driven transcriptomic perturbations (31). They may also explain the recent finding that the avirulent phenotype of *△myr1* parasites during *in vivo* mouse infections is rescuable by co-infecting animals with both wild type and MYR1-deficient parasites (46); parasites expressing MYR1 may induce host cells to secrete paracrine factors that suppress transcription of inflammatory gene products that would otherwise limit *△myr1* parasite infections. Note that while this manuscript was in preparation, a transcriptomic study of U-I macrophages was published by Hunter and colleagues (44). While they used different strains (Pru and CEP), looked at later time points (20-24 hpi), and did not look at MYR1-dependent effects, their primary conclusions that U-I cells experience a major impact of rhoptry effectors (in their case, specifically ROP16), and that paracrine effects are also in play, are similar to the conclusions reached here.

Of note, the host response to MIGs + Parasites was especially subtle, given that parasite invasion involves dramatic mechanical perturbations to host cells that might be expected to trigger transcriptional responses. As we did not use host cells containing MIGs and no parasites or vice versa, we could not discern the impact of the MIGs in isolation. Nonetheless, it is tempting to speculate that the response to MIGs + Parasites may reflect MIGs counteracting the effects associated with parasite penetration and PVM formation, which may explain the subtle net inflammatory response.

In addition, the presence of ROPs and MDGs that respectively activate and suppress certain host processes might be interpreted as energetically wasteful. Why might *Toxoplasma*, an obligate intracellular organism likely under selective pressure for transcriptional efficiency, expend extra resources on effectors that negate one other’s effects? One possibility is that ROPs may have undergone selection optimizing for functions required to establish and protect the parasite’s intracellular niche (4-9, 47-52), but may also trigger unavoidable host responses detrimental to the parasite. In this scenario, MDGs and paracrine effects could ameliorate such ROP-triggered side effects while theoretically leaving processes beneficial to the parasite intact. Another possibility is that ROPs and GRAs could grant *Toxoplasma* the ability to fine tune host responses in terms of timing and/or magnitude, likely an advantage to an organism that must be equipped to encounter a diversity of intracellular environments due to its extraordinarily broad host range (1, 53, 54).

Our analysis also reveals a striking effect on the host cell cycle, in which U-I cells exhibit enrichment of G2/M phase cells. Curiously, this enrichment appeared not to be preserved at 3 hpi, which suggests that the responsible parasite factors exert only a transient influence on host cell cycle-related genes. Of the parasite effectors currently known to modulate the host cell cycle (19, 22, 23, 40, 55), ROP16 is most likely to explain these results: ROP16 phosphorylates ubiquitin-like containing PHD and RING finger domain 1 (UHRF1) in a manner that peaks at 3 hpi, which leads to epigenetic silencing of cyclin B1 (40), a component of the cyclin B1/Cdk1 complex required for the G2/M transition. However, ROP16 is likely not the only ROP to impinge on the host cell cycle, as comparison of G1 phase U-I vs. U-U cells also revealed enrichment in the p53 Pathway and KRAS Signaling Up gene sets (Fig. 4; Fig. 9A and Fig. 9B, ROPs), whose corresponding pathways promote cell cycle progression. Further analysis of U-I cells from yet more time points and particularly those not in G1 may shed more light on this interplay.

Finally, our analyses thus far reflect a fraction of the possible uses of our dataset, which includes variables of time, parasite strain, infection status, cell cycle phases, and UMAP populations. For example, our dataset includes reads not only from host cells but also the *Toxoplasma* parasites, rendering the data a co-transcriptomic resource that will likely illuminate novel host-parasite interactions. Furthermore, because of limiting numbers, the analyses described here dealt primarily with G1 phase cells in UMAP population 1 and largely excluded cells in the remaining cell cycle phases and UMAP populations. The raw and processed transcriptomic data for these remaining cells, including the results of differential gene expression analysis and of gene set enrichment analysis, are publicly available (as are the data described thus far) on the Gene Expression Omnibus under accession number GSE145800. Future analyses of such cells may reveal roles for cell cycle phase, host cell type, or other host-dependent processes.

## Materials and Methods

### Cell and Parasite Culture

All *Toxoplasma gondii* strains were maintained by serial passage in human foreskin fibroblasts (HFFs) cultured at 37°C in 5% carbon dioxide (CO_2_) in complete Dulbeco’s Modified Eagle Medium (cDMEM) supplemented with 10% heat-inactivated fetal bovine serum (FBS), 2 mM L-glutamine, 100 U/ml penicillin, and 100 μg/ml streptomycin.

### Construction of Parasite Strains

RH Tfn-BLA TdTomato parasites were constructed by transfecting ∼10^7^ RH Toxofilin-HA-beta-lactamase (RH Tfn-BLA) parasites (8) with the plasmid pSAT1::Cas9-U6::sgUPRT (56) and with the linearized pCTR_2T_ plasmid containing the construct to express TdTomato (57) using the AMAXA Nucleofector 4D system (U-033 setting) and the P3 primary cell 4D-nucleofector X kit with the 16-well nucleocuvette strip (Lonza, V4XP-3032). Clones were obtained by FACS using the FACS Aria II sorter at the Stanford Shared FACS Facility for the brightest red parasites and single cloning the TdTomato+ enriched population by limiting dilution into 96-well plates.

A CRISPR-Cas9 strategy was used to construct RH *△myr1* mCherry Tfn-BLA parasites from an RH *△myr1* mCherry parental strain (25). The parental strain was transfected with the plasmid pSAG1::Cas9-U6::sgUPRT and a linear construct that contained Toxofilin-HA-Beta-lactamase (Tfn-HA-BLA) expressed under *toxofilin*’s endogenous promoter. The Tfn-HA-BLA construct was PCR amplified from the plasmid SP3 (8) such that the final amplicon was flanked by 20 nucleotide (nt) homology arms to the *UPRT* gene that are identical to those used to target constructs to the *UPRT* locus in (56). 15 μg of pSAG1::Cas9-U6::sgUPRT and 3 μg of the Tfn-HA-BLA linear amplicon were transfected into ∼10^7^ parental strain parasites using the AMAXA Nucleofector 4D system as described above, except the setting on the nucleofector (T-cell human unstimulated HE setting). Selection for parasites with 5 μM 5-fluorodeoxyuridine (FUDR) began at 1 day post-transfection and proceeded for 3 lytic cycles in monolayers of human foreskin fibroblasts (HFFs). The transfected, selected parasite populations were subjected to two rounds of single cloning, one to generate populations of parasites enriched for the presence of the Tfn-HA-BLA construct, and one to purify for individual parasite clones containing the construct, where the readout for the presence of the construct was cleavage of the BLA-cleavable FRET-based dye CCF2-AM, which results in a fluorescence color shift.

IFA of monolayers infected with newly constructed RH *△myr1* mCherry Tfn-BLA parasites was used to validate correct localization of the Tfn-HA-BLA construct to the parasite rhoptry organelles and the absence of expression of host factors known to be induced in the host nucleus as a result of MYR-dependent GRAs. Coverslips seeded with confluent HFF monolayers were infected with putative RH *△myr1* mCherry Tfn-BLA clones for ∼24 hours, fixed in 4% formaldehyde for 20 minutes, permeabilized in 0.2% Triton X-100 for 20 minutes, blocked in 1x PBS containing 3% bovine serum albumin (3% BSA solution) for 1 hour at room temperature, stained with primary and secondary antibodies in 3% BSA solution, and mounted onto glass slides using DAPI-containing VectaShield (Vector Laboratories, H-1200) and sealed with colorless nail polish. For Tfn-HA-BLA localization, primary antibodies were mouse-anti-ROP2/3/4 (1:250 dilution, 4A7, (58)) and rat anti-HA (1:500, clone 3F10, Sigma Aldrich, 11867431001), and secondary antibodies were goat anti-mouse IgG-Alexa 647 (1:2000 dilution, Thermo Fisher, A-21235) and goat anti-rat IgG-Alexa 488 (1:2000 dilution, Thermo Fisher, A-11006). To verify the absence of the MYR1 protein, the primary antibody rabbit anti-c-myc (1:600 dilution, Sigma, M5546) and the secondary antibody goat anti-rabbit IgG-Alexa 594 (1:2000 dilution, Thermo Fisher, A-11012) were used, and coverslips were seeded with either >3-week-old confluent HFF monolayers or with younger monolayers that had been serum starved for at least 24 hours (using 0.5% FBS instead of 10% FBS in the culture medium) to ensure no spurious induction of c-myc in uninfected host cells. Only clones where the anti-HA and anti-ROP2/3/4 signal colocalized and where no expression of c-myc was detected in the infected host nucleus were selected. RH Tfn-HA-BLA TdTomato parasites (32) and the parental RH *△myr1* mCherry parasites were also subjected to both protocols as positive and negative controls, respectively. Coverslips were imaged either on the Stanford Neuroscience Imaging Service core’s Zeiss LSM 710 confocal microscope or on an Olympus BX60 upright fluorescence microscope.

Western blot was used to verify for the correct size of the Tfn-HA-BLA fusion protein expressed in RH *△myr1* mCherry Tfn-BLA parasite clones. Lysates were generated by treatment of parasite pellets with SDS-PAGE loading dye containing 10% beta-mercaptoethanol. The lysates were separated by SDS-PAGE and transferred to a PVDF membrane, and the membrane was blocked with TBST (TBS, 0.05% Tween-20) containing 5% milk for 1.25 hours. The membrane was incubated with horseradish peroxidase (HRP)-conjugated rat anti-HA (clone 3F10) monoclonal antibodies (Roche, Indianapolis, IN) at a dilution of 1:5,000 for 2 hours, and with 1:10,000 dilution rabbit anti-SAG1 for 30 minutes followed by 1:20,000 dilution goat anti-rabbit-HRP for 30 minutes, and developed using the ECL Prime Western Blotting System (Sigma-Aldrich, RPN2232).

### Determination of Time to Division for Infected 10 T1/2 Cells

To determine the time post-infection at which infected 10 T1/2 cells divided, 10 T1/2 host cell monolayers were infected with RH Tfn-BLA TdTomato parasites, and the infected monolayers were imaged by time lapse microscopy.

To prepare the parasites for the infection, the parasites were released from heavily infected monolayers of HFFs by mechanical disruption of the monolayers using sterile, disposable scrapers and passage at least 6 times through a 25 gauge syringe. The parasites were washed by pelleting out HFF debris (133.5-208.5 x g for 5 minutes) and resuspending the parasite pellet generated by spinning the remaining supernatant 469.2 x g in phenol red-negative low serum DMEM.

To prepare the host cell monolayers, 10 T1/2 cells approximately 1 week from their date of thaw were seeded into 12-well tissue culture plates at approximately 6.0 x 10^4^, 1.2 x 10^5^, and 2.4 x 10^5^ cells per well. The 10 T1/2 cells were then incubated for >2 hours in cDMEM, serum starved by incubation at 37°C in 5% CO_2_ in low serum DMEM (i.e., cDMEM containing 0.5% FBS instead of 10% FBS) for 23 hours, washed and stained with 500 μl of Cell Tracker Green CMFDA (CTG-CMFDA; Thermo Fisher, C2925) diluted 1:1000 in prewarmed PBS for 30 minutes at 37°C in 5% CO_2_, and washed and incubated in phenol red-negative low serum DMEM (Thermo Fisher, 31053028) for 30 minutes. At 24 hours post-infection, RH Tfn-BLA TdTomato parasites were added to the monolayers at a multiplicity of infection (MOI) of 6.

Time lapse, epifluorescence images of the infected monolayers were acquired over 16 hours in a controlled (37°C and 5% CO_2_) environment using a Nikon Eclipse inverted microscope (Julie Theriot lab). Images were acquired every 20 minutes using 100 ms of exposure at 25% power for the Phase, mCherry (to visualize parasites), and GFP (to visualize host cytoplasmic CTG-CMFDA) channels. Cells that were uninfected at the start of the time lapse were followed to determine whether they were infected by the end of the time lapse, as determined by: 1) clearing of cytoplasmic CTG-CMFDA in the exact position of the parasite, 2) disappearance of birefringence in the Phase channel upon parasite invasion, and c) the parasite tracking with the cell at all time points following presumed infection. For each cell for which the precise moment of infection was captured in the live video footage, the time to division was determined using the time the cell was infected as the start time and the time the cell divided (if applicable) as the end time.

### FACS of Single Uninfected-Injected (U-I) and Control Cells for Single Cell RNA

#### Sequencing (scRNA-seq)

##### Preparation of Single Cells for FACS Sorting

To generate U-I cells for FACS sorting, 10 T1/2 host cells approximately 1 week from their date of thaw were seeded into 6-well tissue culture plates at a density of approximately 2.6 x 10^5^ cells per well, incubated for >2 hours in cDMEM, and serum starved by incubation in low serum DMEM (i.e., cDMEM containing 0.5% FBS instead of 10% FBS) for 24 hours. 10 T1/2 host cells were chosen as they are monolayer-forming, contact-inhibited fibroblasts that are also suitable for single cell sorting due to adequate dissociation into individual cells by a combination of mechanical and chemical disruption. Next, either RH Tfn-BLA TdTomato or RH *△myr1* mCherry Tfn-BLA parasites were released from heavily infected monolayers of HFFs by mechanical disruption of the monolayers using sterile, disposable scrapers and passage at least 6 times through a 25 gauge syringe. Parasite-free lysate was similarly generated by mechanical disruption of uninfected HFFs. Parasites and parasite-free lysate were washed by pelleting out HFF debris (133.5-208.5 x g for 5 minutes) and resuspending the parasite pellet generated by spinning the remaining supernatant 469.2 x g in low serum DMEM containing either 1 μM DMSO (for the wild type and *Δmyr1* conditions) or with 1 μM of the invasion inhibitor cytochalasin D (cytD, for the cytD-treated wild type condition), incubated at room temperature for 10 minutes, and applied to the serum starved 10 T1/2 monolayers at multiplicity of infection (MOI) = 6, which maximized the abundance of U-I cells in tissue culture. All 10 T1/2 monolayers were spun at 469.2 x g for 5 minutes to synchronize parasite contact with the monolayer. Infections were allowed to proceed for 30 minutes, 1.5 hours, or 2.5 hours at 37°C. Because cytD is a reversible inhibitor, extra cytD-containing low serum DMEM was added to each of the drug-treated infections to maintain the concentration of cytD at 0.5-1 μM for the entire infection duration.

To identify host cells injected by parasite proteins, a 6x stock solution of the beta-lactamase substrate CCF2-AM (Thermo Fisher, K1032) was added to the media over 10 T1/2 cells so that the final concentration of CCF2-AM was 1x. CCF2-AM treated monolayers were incubated under foil (to protect from light) for 30 minutes at room temperature (to prevent breakdown of CCF2-AM, which degrades at 37 °C), bringing up the total infection duration to 1 h, 2 h, and 3 h.

To harvest the 10 T1/2 monolayers for subsequent FACS analysis, the monolayers were: 1) washed 3 times in 1x phosphate buffered saline (PBS) to remove any HFF debris adhering to the monolayer; 2) incubated in trypsin (prepared in plastic vessels only) at room temperature for 6-10 minutes; 3) quenched in an equal volume of FACS buffer (1 x PBS + 2% FBS + 50 mM MgCl_2_*6H_2_0 + 50 μg/ml DNase I); 4) passed 3 times through an 18 gauge syringe to break any residual cell clumps; and 5) washed in FACS buffer to remove excess trypsin. Of note, DNase I and MgCl_2_ were included in the FACS buffer to prevent clumping of cells from cell death. The cells were then stained with a viability dye and an extracellular parasite stain by 1) resuspending them in 500 μl of chilled 4°C 1x PBS containing 3% bovine serum albumin (BSA) and 1:500 dilution rabbit anti-SAG2A primary antibody (gift of C. Lekutis) for 30 minutes on wet ice, 2) washing them in 5 ml of ice cold 1x PBS and spinning at 133.5 x g (lowest setting) at 4°C for 5 minutes, and 3) resuspending the pellets in chilled 1x PBS containing 3% BSA, 1:1000 dilution goat anti-rabbit IgG-Alexa 647 (Thermo Fisher), and 3 μl/ml near-infrared live/dead fixable viability dye (Thermo Fisher, L94375) for 30 minutes on wet ice. Samples were then washed as before, the pellets were resuspended in 1 ml of chilled FACS buffer, and the cell suspension was transferred through a nylon filter cap (Thomas Scientific, 4620F40) into polypropylene FACS tubes stored on wet ice and protected from light until FACS sorting.

##### FACS of Single Cells

To prepare the multi-well lysis plates into which cells were deposited during FACS sorting, lysis buffer was dispensed either by the Mantis liquid handling robot (Formulatrix) at 0.4 μl per well into 384-well hard shell low profile PCR plates (Bio-rad) for single cell RNA sequencing, or by hand at 5 μl per well into 96-well hard shell low profile PCR plates (Bio-rad) for bulk RNA sequencing. Lysis buffer was prepared in batches of 8 ml by mixing 5.888 ml water, 160 μl recombinant RNase inhibitor (Takara Clonetech), 1.6 ml of 10 mM dNTP (Thermo Fisher), 160 μl of 100 μM oligo-dT (iDT), 1:600,000 diluted ERCC spike-in RNA molecules (Thermo Fisher), and 32 μl 10% Triton X-100. All reagents were declared RNase free. Lysis plates were prepared the night before each FACS sort, stored overnight at −80°C, and kept on dry ice during the FACS sort.

All host cell samples were sorted at the Stanford Shared FACS Facility (SSFF) by the BD Influx Special Order sorter using the following channels: forward scatter (488 nm blue laser, SSC detector), side scatter (488 nm blue laser, FSC detector), BV421 (405 nm violet laser, V460 detector, which detected cleaved CCF2-AM), BV510 (405 nm violet laser, V520 detector, which detected uncleaved CCF2-AM), mCherry (561 nm yellow laser, Y610 detector, which detected parasite-associated cells), APC (640 nm red laser, R670 detector, which detected the extracellular parasite stain vs. anti-SAG2A), and APC-Cy7 (640 nm red laser, R750 detector, which detected dead cells). Gating strategy used to obtain U-I, I-I, and U-U cells is indicated in Fig. 2B. More specifically, cells without red fluorescence (i.e., parasite-free) but with enhanced signal from cleaved CCF2-AM (i.e., injected) were sorted as U-I cells, while those with red fluorescence (i.e., parasite-associated), enhanced signal from CCF2-AM (i.e., injected), and low extracellular parasite stain were sorted as I-I cells. Of note, the parasite-associated gate, from which I-I cells were obtained, was intentionally kept narrow to ensure host cells were each infected with approximately one parasite apiece, which limited confounding downstream analysis with penetration of >1 parasite at two different time points. Single color and colorless controls were used for compensation and adjustment of channel voltages. Fluorescence data were collected with FACSDiva software and analyzed with Flowjo software. Cells were index sorted such that each cell’s fluorescence data were recorded for subsequent analysis. For single cell experiments, cells were sorted into the 384-well lysis plates at 1 cell per well. For bulk experiments, cells were sorted into 96-well lysis plates at 50-100 cells per well. Plates were sealed with foil plate sealers and immediately placed on dry ice until the completion of the sort. Plates were then stored at −80°C until library preparation.

### cDNA Synthesis from Single Cell RNA, Library Preparation, and Sequencing

To convert the RNA obtained from single cells and bulk samples to cDNA, we employed the Smart-seq2 protocol (59). For single cell library preparation, the liquid handling robots Mantis (Formulatrix) and Mosquito (TTP Labtech) were employed to transfer and dispense small volumes of reagents, and final reaction volume was 2 μl per well. For bulk sample library preparation, liquid handling was performed with standard multichannel pipets and final reaction volume was 25 μl per sample. cDNA was subject to 19 round of pre-amplification and then quantified using EvaGreen and diluted in EB buffer to obtain a final concentration of 0.4-0.8 ng/μl per sample. Library preparation continued using in-house Tn5 tagmentation. For single cell libraries, we used custom barcoded indices for each cell, and for bulk libraries, we used Nextera XT indices.

Libraries were submitted to the Chan Zuckerberg Biohub Genomics Core for sequencing. Single cell libraries were sequenced on the NovaSeq 6000 by 2×150 base pair paired end sequencing aiming at ∼1 million reads per cell. Bulk libraries were sequenced on the NextSeq by 2×150 base pair paired end sequencing at ∼10 million reads per sample.

### Sequencing Alignment

Reads output from sequencing were aligned to a concatenated genome composed of the mouse genome (GRCm build 38) and the GT1 *Toxoplasma gondii* genome (ToxoDB version 36), which is the most complete reference for type I parasite strains such as the RH strains used in this work. Alignment was performed using STAR, and transcript counting was performed by Htseq-count, with standard parameters used for both packages. A custom python script was used to sum transcript counts to yield a final gene count matrix consisting of all sequenced cells and the number of read counts detected for each gene.

### Data Preprocessing

To filter out cells of poor quality from the analysis, we excluded cells based on the following metrics: gene count, total read sum, percentage of reads that mapped to the mouse-*Toxoplasma* concatenated genome, percentage of reads derived from spiked in ERCC standards, and percentage of reads derived from ribosomal RNA.

The gene count matrices were then normalized as counts per median (cpm). Briefly, we first calculated the sum of reads for all cells. We then divided the read counts by the corresponding sum of reads in each cell and multiplied the fractional count by the median of the sum of reads as a scaling factor. Normalized data were transformed to log2 space after adding a pseudo-count of 1 for each gene of each cell.

To determine the detection limit of each experimental trial (e.g., the 50% detection rate), we computed a logistic regression model from a plot of the detection probability for spiked-in ERCC standards. We then excluded genes where less than 5 cells in the experimental trial expressed that gene at a level above the detection limit.

To identify host genes associated with infection, we first excluded mouse genes to which *Toxoplasma* sequences erroneously map (Table S1) by aligning RNA sequences obtained from single cell extracellular RH parasites to the concatenated mouse-*Toxoplasma* genome and eliminating all mouse genes with an average log_2_(cpm+1) expression of 0.2 or greater.

#### Single Cell vs. Bulk Sample Correlation Analysis

To validate the single cell expression data, we plotted the log2 mean expression of each differentially expressed gene, as identified by the MAST algorithm using the single cell data, in single cells vs. bulk samples. The sklearn package was applied to compute linear regression and the corresponding R^2^ (coefficient of determination) values.

#### Cell Cycle Analysis and Annotation

To predict the cell cycle phase of individual single cells, we curated a list of 175 cell cycle marker genes from the literature (60–67) and from the database CycleBase 3.0 (68). We computed the first two principle components of the gene count matrix using principle component analysis (PCA) and projected the cells. We partitioned the cells using K-Means clustering and assigned the clusters with their predicted cell cycle phases (G1, S, and G2/M) based on the expression of 175 cell cycle marker genes curated from the literature (Table S2).

#### Dimensionality Reduction

To visually represent relationships between single cells across all experimental trials based on their transcriptional variation regardless of experimental conditions, we identified and filtered for the top 1000 genes with the highest dispersion (i.e., the genes with the most variable expression for their bin groups with similar expression level) across all datasets, applied mutual nearest neighborhood batch correction (MNNPY) to correct for batch effects, and projected the data onto two dimensional space with the uniform manifold approximation and projection (UMAP) algorithm using default parameters in Scanpy (69). Leiden clustering using the top 1000 dispsered genes enabled partitioning of cells into three populations (1, 2, and 3), which we separated into individual datasets for downstream analysis.

#### Infection Status Classification

We also determined the host infection load by quantifying the percent of reads that mapped to *Toxoplasma* in a given sample. We filtered samples which were shown by FACS to exhibit one presumed infection status (based on red fluorescence from internalized parasites) but were determined to exhibit the opposite or an ambiguous infection status otherwise (based on percentage of reads derived from *Toxoplasma*).

#### Differential Expression Analysis

To obtain differentially expressed genes (DEGs) between all pairs of conditions, we used the Model-based Analysis of Single Cell Transcriptomics (MAST) algorithm (70) to compute the results on all G1, correctly classified, and UMAP population 1 cells. We used the default settings except with an adaptive conditional mean of expression based on 20 number of bins, at least 30 genes in each bin, and we did not filter out any gene with non-zero expression frequency in the samples.

#### Gene Set Enrichment Analysis

We performed gene set enrichment analysis (GSEA) on the lists of differentially expressed genes between all pairs of experimental conditions, ranked by their relative expression in each of the two conditions, for each individual experimental trial using the fast pre-ranked gene set enrichment analysis (fgsea) package (71). Genes were compared to the Molecular Signature Database’s Hallmarks gene sets. Pathways with an adjusted p-value, i.e., false discovery rate (FDR) of < 0.25 were considered to be significantly enriched at the top or the bottom of the ranked list of differentially expressed genes.

### Identification of Differentially Regulated Genes between Conditions Using CCF2-AM

#### Ratio Correlation Analysis

To identify host genes associated with injection, we first computed the CCF2-ratio for each cell, a metric that serves as a readout for parasite effector injection and that was calculated by dividing the log-transformed cleaved CCF2-AM fluorescence by the log-transformed uncleaved CCF2-AM fluorescence. Next, we excluded mouse genes below the detection limit and mouse genes to which *Toxoplasma* sequences erroneously map. Finally, for each infected or injected condition (i.e., Wt U-I, Wt I-I, Dmyr1 U-I, Dmyr1 I-I, CytD U-I, and CytD I-I), we computed the Spearman correlation of each of the remaining genes to the CCF2-AM ratio using cells from the condition of interest and cells from the cognate U-U condition (e.g., to calculate Spearman correlations for Wt U-I cells, we correlated gene expression to the CCF2-AM ratio in Wt U-I and Wt U-U cells).

To identify genes differentially regulated between pairs of conditions using the CCF2-AM correlation data, we generated scatterplots where each data point represented a gene and its displacement on each of the x- and y-axes represented the Spearman correlation in each of the conditions. We modeled a Gaussian distribution using the Sklearn package with default parameters with settings “n_components=1” and “covariance_type=’full’”. Genes with a difference in CCF2-AM correlation score of >0.2 or <-0.2 and whose probability densities were more than three standard deviations from the mean probability density were considered to exhibit significant differential expression between the pair of conditions under scrutiny.

#### Generation of Strip Plots

Strip plots in Fig. 8B were generated using seaborn’s catplot and boxplot packages to plot the normalized expression scores for each cell across 11 experimental conditions for differentially expressed genes (DEGs) between Dmyr1 U-U and Mock samples. The normalized expression score for each cell was calculated by 1) subtracting the minimum log2 cpm for that gene across all cells in the experiment, 2) dividing the difference by the maximum log2 cpm across all cells, such that the cell with the lowest count received a score of 0 and the cell with the highest count received a score of 1, and 3) computing the average of the normalized cpm’s for each cell.

#### Generation of Bubble Plots

Bubble plots in Fig. 9A and Fig. 9B were generated using a custom Python script in which columns indicated the gene set and rows indicated the comparison (taken to signify the impact of one to a few parasite-dependent stimuli on host transcription) from which the gene sets were enriched. The bubbles in each plot were color coded and sized based on one of two schemes. In the first scheme, bubble color indicated the normalized enrichment score, where scores were positive if the genes corresponding to a given gene set were expressed higher in the first member of the pair in the given comparison than in the second member of the pair, and bubble size indicated the significance (i.e., false discovery rate, or FDR) of the enrichment, where the absolute size of each bubble corresponded to the reciprocal of the FDR. In the second scheme, bubble color indicated the log2 normalized fold change in expression of DEGs from a given comparison that also corresponded to a given Hallmark pathway, where positive fold changes indicated genes were expressed higher in the first member of the pair in the comparison than in the second member, and bubble size indicated the number of DEGs used to calculate the fold change. To calculate fold change for each given comparison’s gene set, DEGs from the comparison that fell under the gene set were identified, expression of these DEGs in cpm was normalized across all cells to a maximum value of 1 and averaged across all the cells in each condition, and average normalized expression in the first experimental condition was divided by average normalized expression in the second experimental condition.

#### Data Availability

The RNA sequencing dataset produced in this study has been uploaded in its entirety to the National Center for Biotechnology Information (NCBI) Gene Expression Omnibus (GEO) database under accession number GSE145800. The dataset includes the raw fastq files, processed gene count files (in counts per median), an anndata file containing the processed gene count files and other metadata such as the cell cycle phase and percentage of *Toxoplasma*-derived reads for each cell, and results of differential gene expression analysis and gene set enrichment analysis for up to eleven distinct species of host cell (i.e., Wt U-I, Wt I-I, Wt U-U, Dmyr1 U-I, Dmyr1 I-I, Dmyr1 U-U, CytD U-I, CytD I-I, CytD U-U, Mock, and CytD Mock) at each of three time points (i.e., 1, 2, and 3 hours) post-infection. Here, Wt, Dmyr1, and CytD designate host cells arising from monolayers infected with wild type (RH Tfn-BLA TdTomato), *Δmyr1* (RH *△myr1* mCherry Tfn-BLA) or cytochalasin D-treated wild type parasites, respectively, while Mock and CytD Mock refer to cells arising from monolayers mock-infected with parasite-free lysate (where the lysate was pretreated with cytochalasin D in the CytD Mock condition).

## Acknowledgments

Special thanks to Terence Theisen for his assistance in preparing U-I cells for FACS sorting; Dr. Anita Koshy for her careful reading of the manuscript; Meredith Weglarz and Marty Bigos from the Stanford Shared FACS Facility for their assistance during the FACS sorts for U-I cells; and David Parks and Catherine Carswell-Crumpton for their input in optimizing the FACS sort conditions to obtain pure U-I cell populations. We also thank Saroja Kolluru and Robert Jones for their assistance during library preparation and sample submission; Fabio Zanini for experimental suggestions; Geoff Stanley and Michelle Chen for their input on the bioinformatic analysis of the U-I scRNAseq data; and Julie Theriot, Zhou Xiaoxue, and Matthew J. Footer for their assistance with the live video microscopy experiments. This work was supported by NIH F30 AI124589-03 (SR), Stanford Interdisciplinary Graduate Bio-X Fellowship (YX), RO1 AI021423 and AI129529 (JCB), Chan Zuckerberg Biohub (SRQ), NIH S10 Shared Instrument Grant S110RR025518-01 (Stanford Shared FACS Facility), and NIH NS069375 (Neuroscience Microscopy Core).

## SUPPLEMENTARY FILE FIGURE AND TABLE LEGENDS

**Figure S1.** Construction and validation of RH Δ*myr1* mCherry Toxofilin-HA-beta-lactamase (RH Δ*myr1* mCherry Tfn-HA-BLA). **(A)** CRISPR/Cas9 strategy to construct RH Δ*myr1* mCherry Tfn-HA-BLA by disrupting parasite UPRT (blue) in parental strain RH Δ*myr1* mCherry. The parental strain was transfected with plasmid pSAG1::-Cas9-U6::sgUPRT (pink) and a linear amplicon containing Tfn-HA-BLA (purple) and flanked with 20 nucleotide homology arms to UPRT (blue). The region of UPRT complementary to the guide RNA, which also contains the site at which Cas9 cleaves, is in pink. **(B)** Western blot for the Tfn-HA-BLA construct in RH Δ*myr1* mCherry Tfn-HA-BLA. All lanes were obtained from the same gel. **(C)** Immunofluorescence assay (IFA) for colocalization of the Tfn-HA-BLA protein product and ROP2/3/4. **(D)** FACS analysis of 10 T1/2 cells infected with parasite-free lysate, RH Tfn-HA-BLA, and RH Δ*myr1* mCherry Tfn-HA-BLA. Cleavage of reporter dye CCF2-AM indicates injection of the Tfn-HA-BLA construct.

**Figure S2.** Human foreskin fibroblast (HFF) feeder cell debris contaminates the uninfected-injected (U-I) FACS gate. In all panels, a host cell monolayer is treated with either a parasite-free lysate of HFFs or RH Tfn-HA-BLA parasites syringe released from HFFs and then incubated with CCF2-AM to reveal (un)injected cells. *Left:* Infection with RH Tfn-HA-BLA reveals uninjected and injected cell populations. *Middle:* Mock “infection” with para-site-free lysate reveals HFF feeder cell debris contaminating the “injected” cell population. *Right:* Infection with parasite-free lysate that was washed to remove HFF debris reveals a reduction in contamination of the injected population.

**Figure S3. (A)** Serum starvation for 24 h partially inhibits cell division of 10 T1/2 host cells, reducing the possibility of capturing U-I cells that arise from division of an infected host cell (U-Id cells) rather than from an aborted invasion event. Note that the S phase population in the bottom right panel (serum replete, infected cells) also contains G1 phase cells containing parasites, as parasite nuclear content enhances the propidium iodide signal in these cells. **(B)** Histogram depicting the number of infected host cells that divided at various times post-infection, as determined by live video microscopy footage of 200 serum starved 10 T1/2 cells for which the precise moment of infection was captured on camera. Of the 200 infected 10 T1/2 cells, 53 divided over a 16 hour time course, and none divided at earlier than 3.67 hours post-infection.

**Figure S4. Quality control metrics for single cell RNA sequencing data. (A)** Comparison of gene counts (number of genes for which reads from each cell mapped to the concatenated mouse-*Toxoplasma* genome, y-axis) and read sum (total reads, x-axis) for all experimental trials. Cells that passed quality control are indicated in color. **(B)** Percentages of total reads that mapped to open reading frames (ORFs) in the mouse-*Toxoplasma* concatenated genome. **(C)** *Top panel:* Linear regression modeling of measurement accuracy fitted on ERCC spike-ins with abundance above the detection limit. The text within each subplot denotes the coefficient of determination for the regression fit. *Bottom panel:* Logistic regression modeling of detection limit based on ERCC spike-ins. The 50% detection rate is indicated with a black dotted line, and the text within each subplot indicates the detection limit for each experiment in absolute molecular counts. **(D)** Linear regression fitted to scatterplot of average gene counts of differentially expressed genes for single cell RNA sequencing data (x-axis) vs. bulk RNA sequencing data (y-axis). Each point represents a DEG. Text within each subplot denotes the coefficient of determination (R^2^) for the regression. **(E)** The coefficients of determination for linear regression lines fit to scatterplots as in (D) for all possible combinations of single vs. bulk RNA-seq combinations.

**Figure S5. Single cell resolution enables strategic partitioning of individual cells for downstream analysis. (A)** Percentage of reads derived from *Toxoplasma* validates infection status of individual cells. Cells are scored as uninfected if left of the lower decision line (bold, dashed), infected if right of the upper decision line (dotted), and ambiguous if between the decision lines (cross-hatched section). **(B)** Principal component analysis (PCA) projection of cells based on 175 curated cell cycle markers and subsequent Leiden clustering enables partitioning of cells by predicted cell cycle states, G1 (green), S (gold), and G2/M (purple). **(C)** Proportion of cells from each experimental condition in each cell cycle phase. **(D)** Dimensionality reduction and projection of single cells using the *Uniform Manifold Approximation and Projection* (UMAP) algorithm reveals 3 putative cell populations. All panels are reproduced copies of the same projection, each of which is color coded by specific parameters. *Left:* Louvain clusters, used to assign the cells to populations 1, 2, and 3; *middle:* cell cycle phase; *right (3 panels):* experimental trial (1 hpi, 2 hpi, and 3 hpi).

**Figure S6.** Effect of cytochalasin D treatment on the proportion of 10 T1/2 host cells that are U-I cells. **(A)** U-U and U-I cells that arise from Wt and CytD infections at 2 hpi. **(B)** U-U and U-I cells that arise from Wt and CytD infections at 3 hpi.

**Table S1:** Mouse genes discarded due to reads from extracellular RH parasites mapping to these genes in the concatenated mouse-*Toxoplasma* genome.

**Table S2:** Curated list of genes used to assign cell cycle phases to single host cells.

**Table S3:** Differentially expressed genes (DEGs). In all tables, fdr = false discovery rate, and log2FC is the log_2_ fold change in expression of DEGs between the indicated conditions. **(A)** All differentially expressed genes at all time points. **(B)** Wt I-I vs. Mock Differentially Expressed Genes. **(C)** Wt U-I vs. Wt U-U Differentially Expressed Genes. **(D)** Dmyr1 U-I vs. Dmyr1 U-U Differentially Expressed Genes. **(E)** CytD U-I vs. CytD U-U Differentially Expressed Genes. **(F)** Wt U-I vs. Wt I-I Differentially Expressed Genes. **(G)** Dmyr1 U-I vs. Dmyr1 I-I Differentially Expressed Genes. **(H)** Wt I-I vs. Dmyr1 I-I Differentially Expressed Genes. **(I)** Wt U-U vs. Mock Differentially Expressed Genes. **(J)** Dmyr1 U-U vs. Mock Differentially Expressed Genes. For (F) and (G), a gene is designated as showing evidence of being acted upon by counterbalancing effectors (indicated in the “Evidence of Effectors in Wt I-I Cells Counteracting Effectors in Wt U-I Cells?” and “Evidence of Effectors in Dmyr1 I-I Cells Counteracting Effectors in Dmyr1 U-I Cells?” columns, respectively) if it exhibits either: 1) upregulation in one cell type and downregulation in the other compared to a common Wt U-U standard, or 2) more upregulation or downregulation in Wt U-I cells compared to the standard than in Wt I-I cells.

**Table S4:** CCF2-AM correlation data for **(A)** Wt U-I vs. Wt I-I cells and **(B)** Dmyr1 I-I vs. Wt I-I cells. A gene is designated as showing evidence of being acted upon by counterbalancing effectors (indicated under the “Evidence of Effector Counterbalancing?” column) if it exhibits either: 1) negative correlation with CCF2-AM in one cell type and a positive CCF2-AM correlation in the other cell type, or 2) stronger positive or negative correlations in Wt U-I cells than in Wt I-I cells.

