## Supplementary figures and images for "Differential Impacts on Host Transcription by ROP and GRA Effectors from the Intracellular Parasite *Toxoplasma gondii*"

### Figure S1

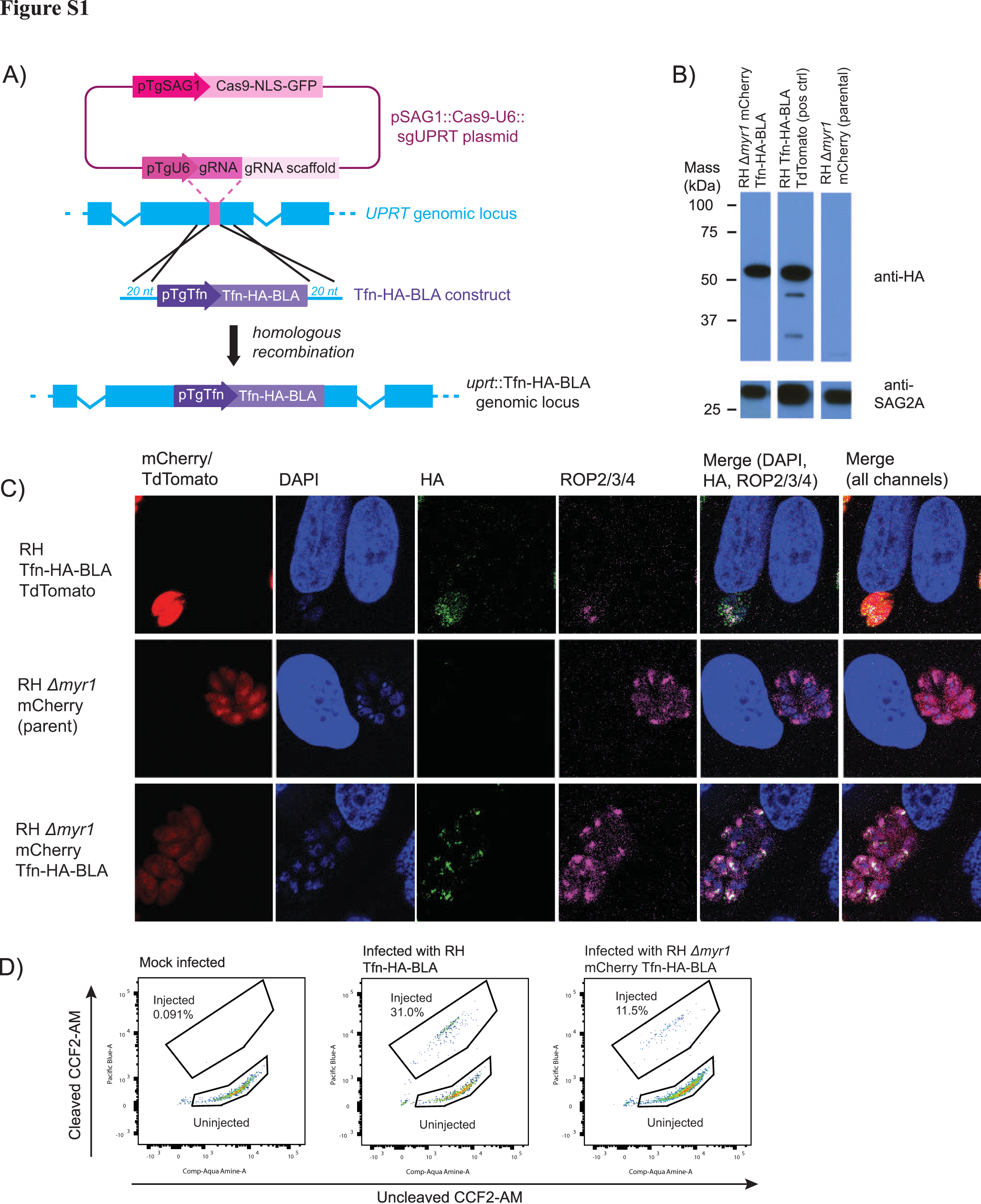

### Figure S2

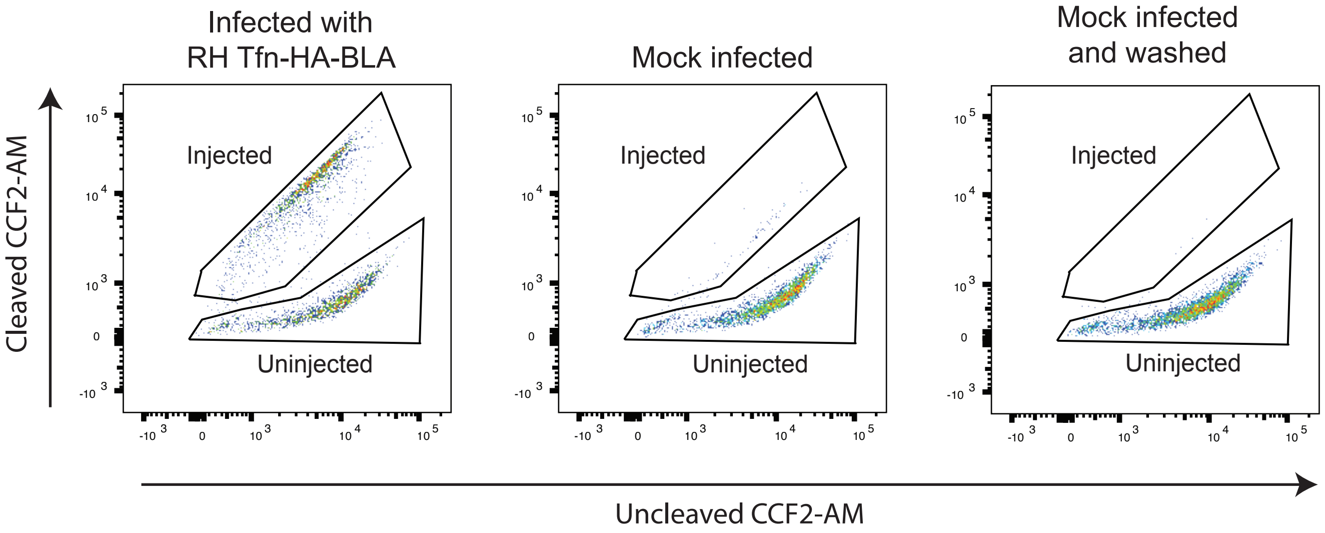

### Figure S3

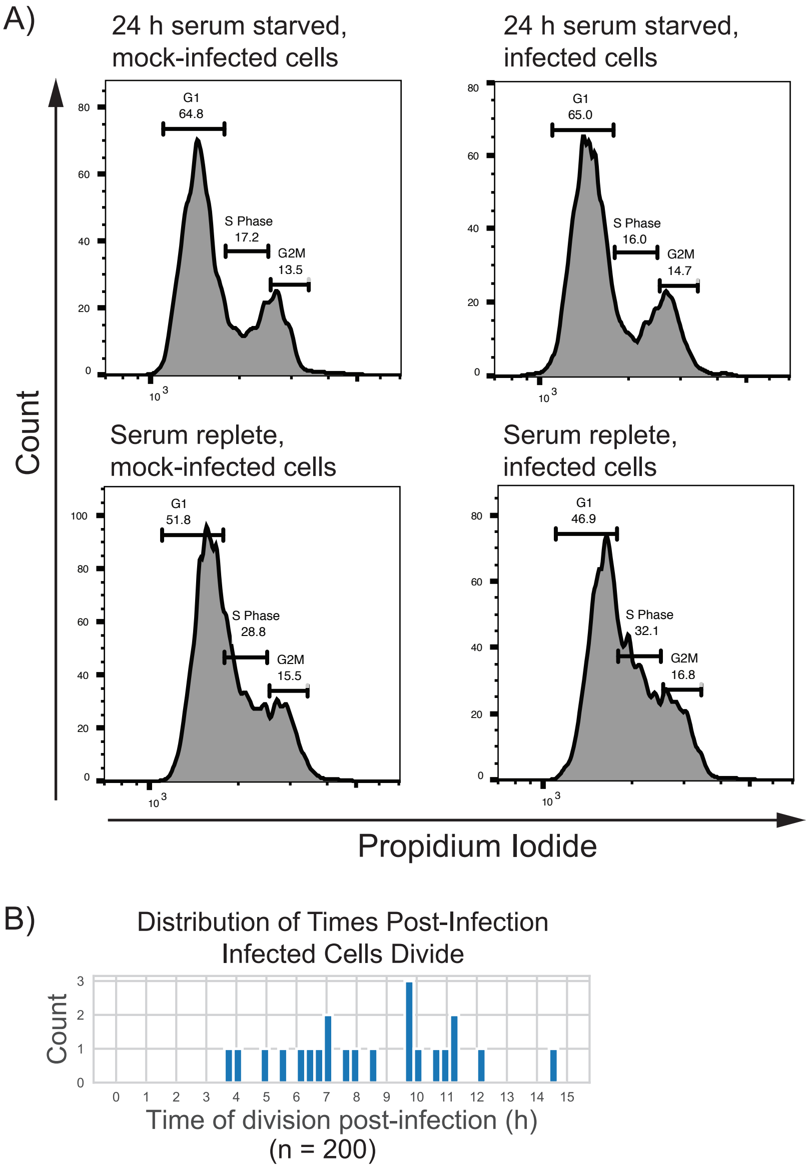

### Figure S4

**Figure S4**

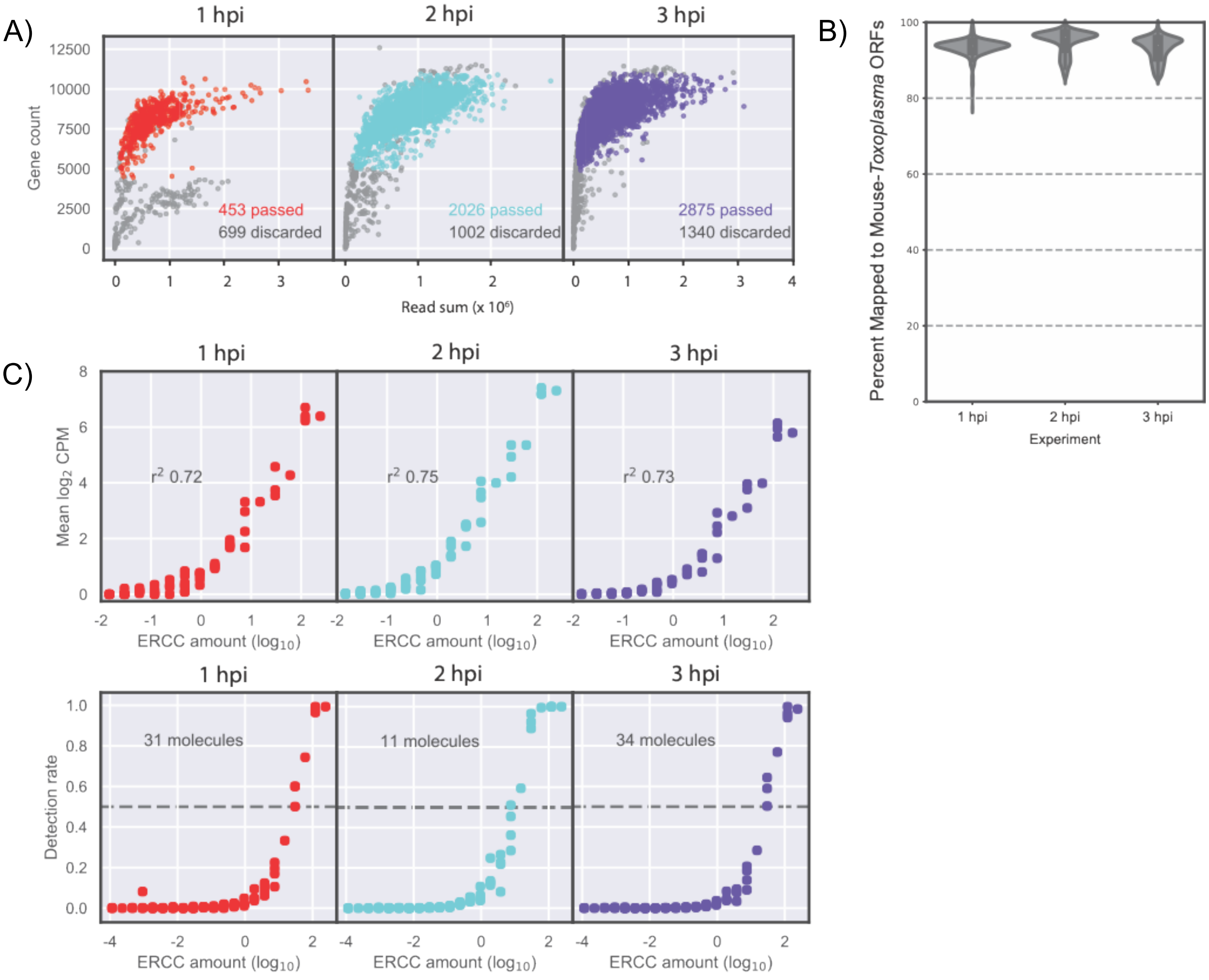


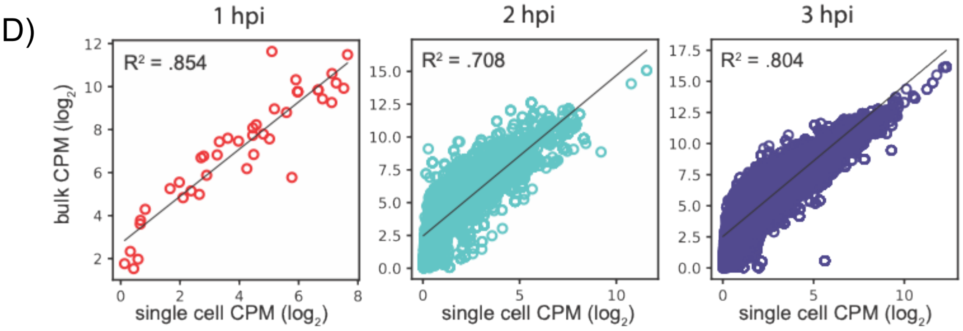


E)

| | Bulk 1 hpi | Bulk 2 hpi | Bulk 3 hpi |
|---|---|---|---|
| Single Cell 1 hpi | 0.854 | 0.655 | 0.630 |
| Single Cell 2 hpi | 0.742 | 0.708 | 0.788 |
| Single Cell 3 hpi | 0.669 | 0.621 | 0.804 |

### Figure S5

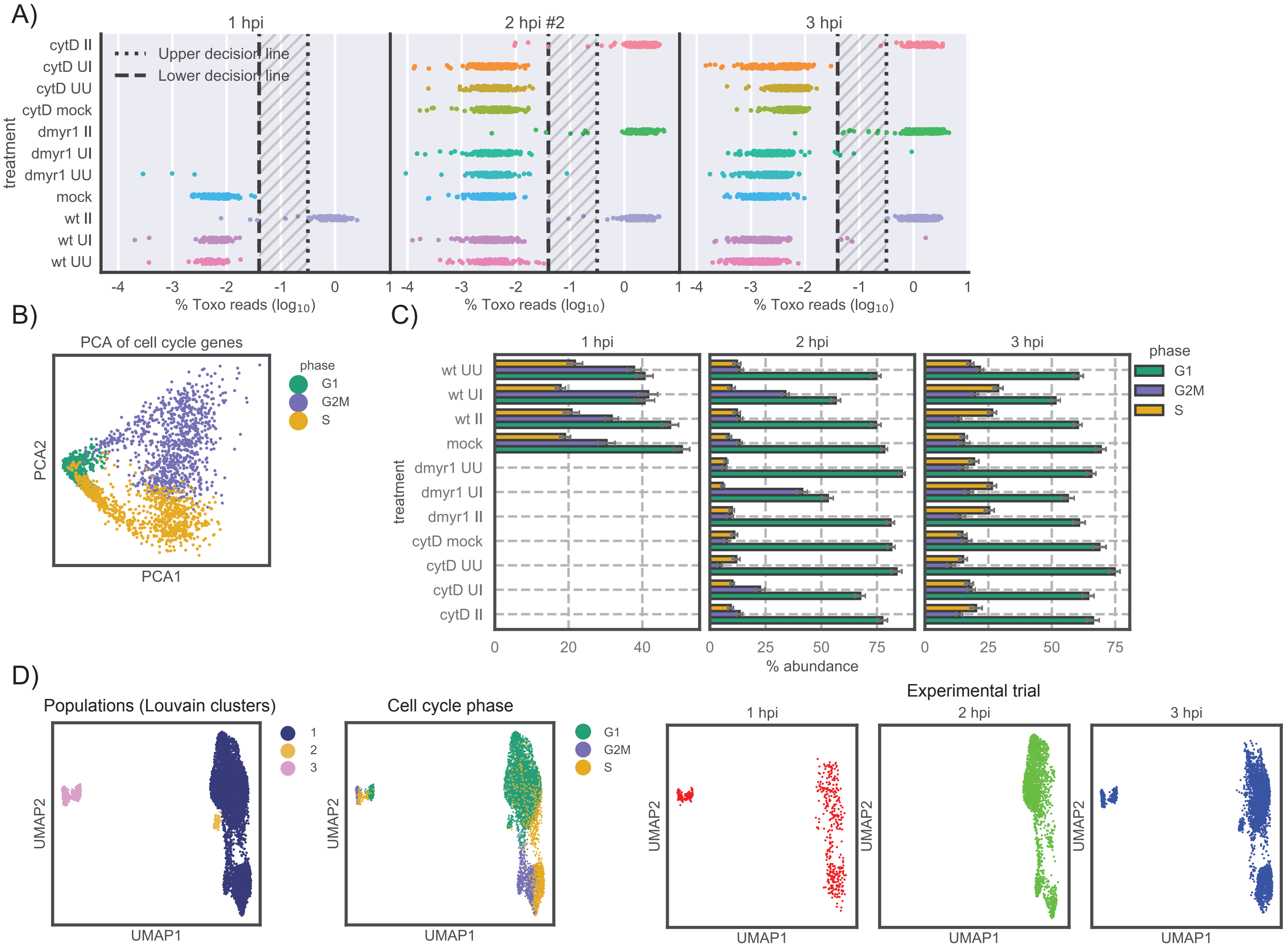

### Figure S6

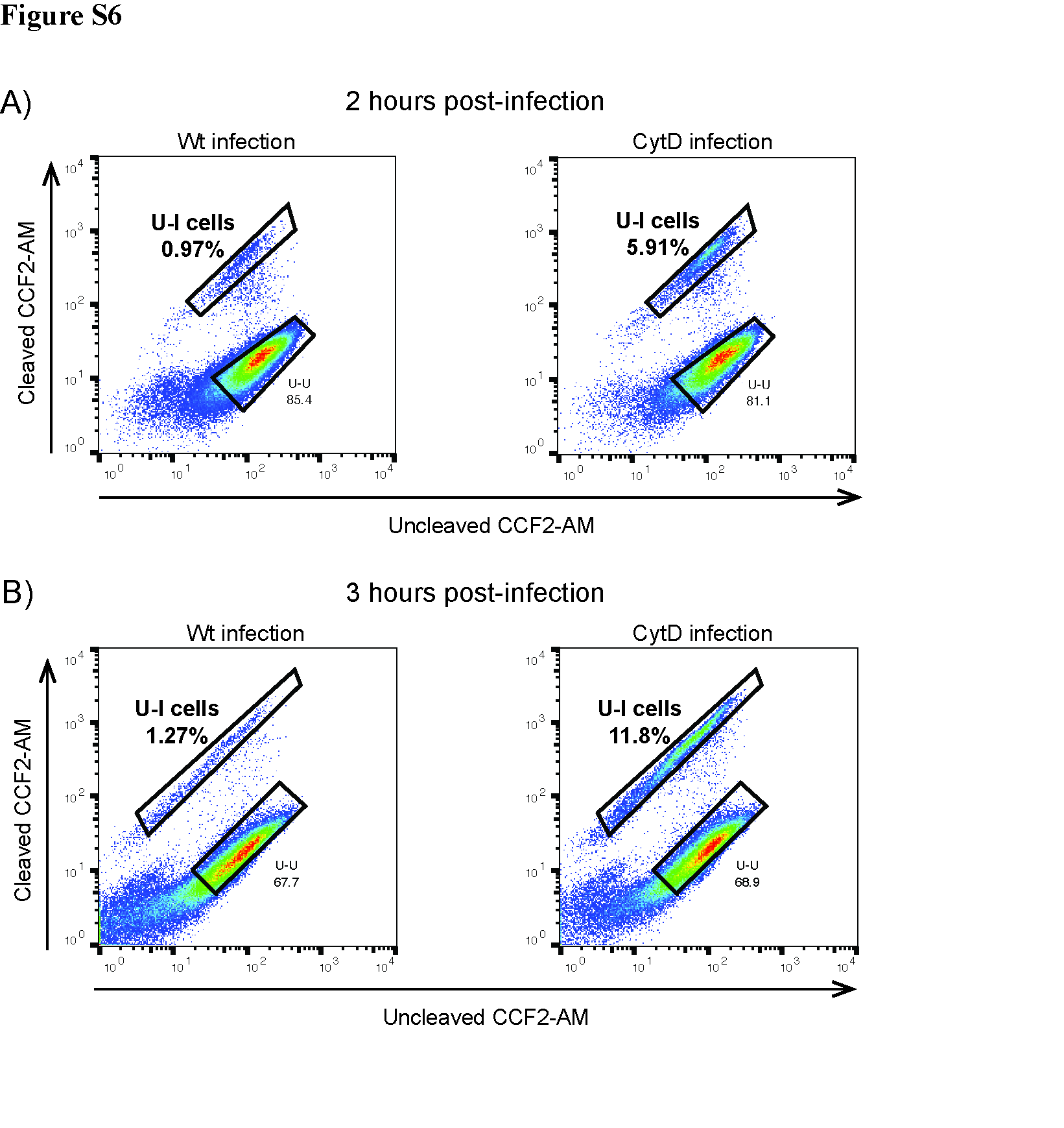
